# Efficient phase coding in hippocampal place cells

**DOI:** 10.1101/630319

**Authors:** Pavithraa Seenivasan, Rishikesh Narayanan

## Abstract

Hippocampal place cells encode space through phase precession, whereby neuronal spike phase progressively advances during place-field traversals. What neural constraints are essential for achieving efficient transfer of information through such phase codes, while concomitantly maintaining signature neuronal excitability specific to individual cell types? We developed a conductance-based model for phase precession in CA1 pyramidal neurons within the temporal sequence compression framework, and defined phase-coding efficiency using information theory. We recruited an unbiased stochastic search strategy to build a model population that exhibited physiologically observed heterogeneities in intrinsic properties. Place-field responses elicited from these models matched signature sub- and supra-threshold place-cell characteristics, including phase precession, sub-threshold voltage ramps, increases in theta-frequency power and firing rate during place-field traversals. Employing this model population, we show that disparate parametric combinations with weak pair-wise correlations resulted in models with similar high-efficiency phase codes and similar excitability characteristics. Mechanistically, the emergence of such parametric degeneracy was dependent on the differential and variable impact of individual ion channels on phase-coding efficiency in different models, and importantly, on synergistic interactions between synaptic and intrinsic properties. Furthermore, our analyses predicted a dominant role for calcium-activated potassium channels in regulating phase precession and coding efficiency. Finally, change in afferent statistics, manifesting as input asymmetry, induced an adaptive shift in the phase code that preserved its efficiency, apart from introducing asymmetry in sub-threshold ramps and firing profiles during place-field traversals. Our study postulates degeneracy as a potential framework to attain the twin goals of efficient temporal coding and robust homeostasis.

**SIGNIFICANCE STATEMENT:** Neuronal intrinsic properties exhibit significant baseline heterogeneities, and change with activity-dependent plasticity and neuromodulation. How do hippocampal neurons encode spatial locations through the precise timings of their action potentials in the face of such heterogeneities? Here, employing a unifying synthesis of the temporal sequence compression, efficient coding and degeneracy frameworks, we show that there are several disparate routes for neurons to achieve high-efficiency spatial information transfer through such temporal codes. These disparate routes were consequent to the ability of neurons to produce precise encoding through distinct structural components, critically involving synergistic interactions between intrinsic and synaptic properties. Our results point to an explosion in the degrees of freedom available to a neuron in concomitantly achieving efficient coding and excitability homeostasis.

---

Hippocampal place cells encode space using both rate and temporal codes (Fig. 1,*A*). Whereas a rate code is defined by an increase in the firing rate of a neuron within the place field, the associated precession of neuronal spike phase relative to the extracellular theta rhythm characterizes a phase code. The phase code, unlike the non-monotonic rate code where the firing rate peaks around the place-field center, manifests as a relatively monotonic dependence of the neuronal firing phase on the animal’s spatial location *within* the place field (O’Keefe and Dostrovsky, 1971; O’Keefe, 1976; O’Keefe and Recce, 1993; Buzsaki and Moser, 2013; Moser et al., 2015). Such monotonicity confers upon the phase code an enhanced potential for information transfer, with the ability to act as a fine-grained spatial code within a single place field. Although the existence of the phase code within a single place field has been well studied (O’Keefe and Dostrovsky, 1971; O’Keefe, 1976; O’Keefe and Recce, 1993; Skaggs et al., 1996; Mehta et al., 2002; Dragoi and Buzsaki, 2006; Harvey et al., 2009; Schmidt et al., 2009; Geisler et al., 2010; Buzsaki and Moser, 2013; Moser et al., 2015), the efficiency of such a phase code in representing spatial information or the neural constraints that are essential for achieving such efficiency have not been assessed. Specifically, how do hippocampal neurons utilize the finite phase span (0–360**°**) available for efficiently encoding space, involving the statistics of afferent inputs onto a single neuron during place-field traversals? How to define and quantify the efficiency of the phase code with reference to a single neuron encoding a single place field? What roles do neuronal intrinsic properties, individual ion channel properties, and associated heterogeneities within the specific neuronal subtype (Cembrowski and Spruston, 2019) play in characterizing the phase code, and in efficiently utilizing the finite phase span to encode space? Are there specific ion channel combinations and specific intrinsic properties that are essential to the emergence of an efficient phase code *concomitant* with the maintenance of signature electrophysiological properties of hippocampal neurons? How do changes in the statistics of place field-driven afferent inputs onto a single neuron alter the phase code towards preserving coding efficiency?

**Figure 1.**
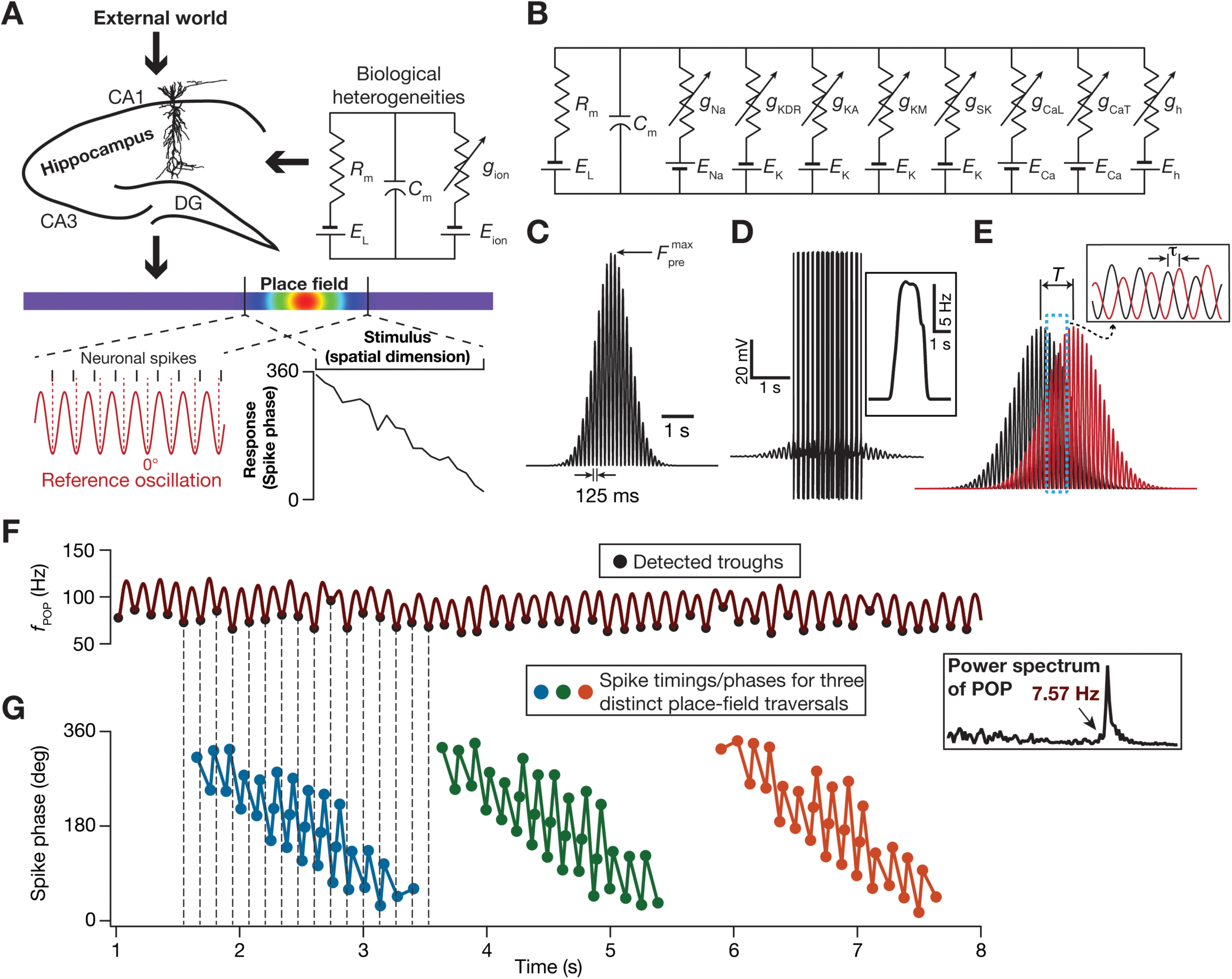
Developing a conductance-based neuronal model that incorporates biological heterogeneities to assess efficiency of the phase code within the temporal sequence compression framework. (A) Hippocampal rate and phase codes: A representation of hippocampal neurons, endowed with inherent biological heterogeneities in active and passive neuronal properties, receiving dynamic spatial stimuli from the external world. The rate code (violet–red along the rainbow spanning lower to higher firing) during an animal’s traversal along a one-dimensional track corresponds to a bell-shaped profile in the neuronal firing rate within the place field of the neuron. The concurrent phase code is derived from the phase of neuronal spikes with respect to an external reference oscillation. (B) Electric circuit equivalent of the conductance-based neuronal model employed in this study. (C–D) A Gaussian modulated sinusoid (C) defined the probability distribution of activating 100 independent synaptic inputs arriving onto the conductance-based model during a place field traversal. In response to afferent synaptic activation, the neuronal model elicited a voltage response (D), with spike rate defining the rate code (D; inset). (E) Illustration of the temporal relationship between the probability distributions that govern the two adjacent place field afferent synaptic activation. *T* signifies the longer time scale that corresponds to the temporal distance (the travel time) between adjacent place field centers. *τ* characterizes the shorter theta time scale temporal difference between adjacent place fields, modeled as a phase shift in adjacent sinusoids at theta frequency (inset). For this illustration, *T*=1000 ms; *τ* =75 ms. (F–G) 50 overlapping place field inputs, constrained by the TSC framework, were presented to the model and the cumulative firing rate (*f*_POP_) spanning all such presentations were computed (F). The oscillatory frequency of this cumulative firing rate was computed from its Fourier spectrum (F; inset). Phase coding emerges in the model as a precession of the phase of spikes elicited during each place field traversal (3 traversals shown), computed with *f*_POP_ as the reference oscillation (G).

To address these questions, we built a conductance-based model for phase precession in CA1 pyramidal neurons involving a synthesis of the temporal sequence compression (TSC) framework (Skaggs et al., 1996; Dragoi and Buzsaki, 2006; Geisler et al., 2010), information-maximization approaches for defining efficient codes (Shannon, 1948; Barlow, 1961; Bell and Sejnowski, 1995, 1997; Stemmler and Koch, 1999; Fairhall et al., 2001; Simoncelli and Olshausen, 2001; Lewicki, 2002; Simoncelli, 2003) and the degeneracy framework where similar function could be achieved through disparate structural components (Edelman and Gally, 2001; Prinz et al., 2004; Marder and Taylor, 2011; Rathour and Narayanan, 2019). Employing an unbiased stochastic search involving thousands of heterogeneous conductance-based models, we arrive at a model population that matched electrophysiological signatures of CA1 pyramidal neurons. Using this model population, we show that efficient transfer of spatial information through spike phases could be achieved through multiple disparate routes while concomitantly maintaining signature excitability properties. Mechanistically, our analyses showed that there was no strong dependence of coding efficiency on either intrinsic excitability or the overall afferent synaptic drive. Instead, we found phase-coding efficiency to be an emergent property that is driven by synergistic interactions between synaptic and intrinsic properties. Further, employing the virtual knockout framework, we showed that the impact of individual ion channels on phase-code efficiency was differential and variable. Finally, by modifying the statistics of the afferent inputs within the TSC framework, we demonstrate that asymmetry in place-field afferent inputs introduces predictive temporal shifts to the rate and phase codes, with the change in the phase code constituting an adaptive shift to preserve efficiency. Together, our study derives a clear definition of efficient phase coding and unveils degeneracy in achieving efficient phase coding and signature excitability characteristics.

## METHODS

A principal question addressed in this study pertains to the specific roles of neuronal intrinsic properties and associated ion channels in regulating phase codes of space and the efficiency of such codes in transferring spatial information. An implicit requirement for addressing these questions is a conductance-based model for phase precession that incorporates neuronal biophysical properties, and ion channel heterogeneities (Fig. 1*A*).

### Single compartmental conductance-based model: Passive, active and synaptic properties

We constructed a single compartmental cylinder of 110-µm diameter (*d*) and 97-µm length (*L*). The passive properties of the cylinder were: specific membrane resistance, *R*_m_=40 kΩ.cm^2^ and specific membrane capacitance, *C*_m_=1 µF/cm^2^. The geometric characteristics and the *R*_m_ were chosen such that the *passive* input resistance of the model (=*R*_m_/(π*dL*)=119.3 MΩ) matched with electrophysiological values of ∼120 MΩ, and the *passive* charging time constant (=*R*_m_*C*_m_=40 ms) was ∼40 ms (Narayanan and Johnston, 2007, 2008). The active properties included 8 active ion channels (a total of 9 channels, including the passive leak channel): fast Na^+^ (NaF), delayed rectifier K^+^ (KDR), *A*-type K^+^ (KA), *L*-type Ca^2+^ (CaL), calcium gated K^+^ (SK), hyperpolarization activated cyclic nucleotide gated (HCN), *M*-type K^+^ (KM) and *T*-type Ca^2+^ (CaT) channels. The kinetic schemes for these channels were derived from electrophysiological recordings from CA1 pyramidal neurons: fast Na^+^, CaL, KDR and KA (Magee and Johnston, 1995; Hoffman et al., 1997; Migliore et al., 1999), HCN (Magee, 1998; Poolos et al., 2002), CaT (Shah et al., 2011), SK (Sah and Isaacson, 1995; Sah and Clements, 1999) and KM (Migliore et al., 2006), and the overall voltage dynamics evolved as (Fig. 1*B*):

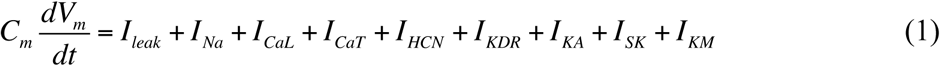

All channels except the SK channel were modeled using the Hodgkin-Huxley formulation; SK channels were modeled using a six-state kinetic model (Sah and Isaacson, 1995; Sah and Clements, 1999). Currents through the sodium channel, the HCN channel and all potassium channels were modeled using the Ohmic formulation, but calcium channels were modeled using the Goldman–Hodgkin–Katz (GHK) formulation (Goldman, 1943; Hodgkin and Katz, 1949), to account for the large concentration gradient observed in the calcium ion. The maximal conductances associated with the individual ionic currents 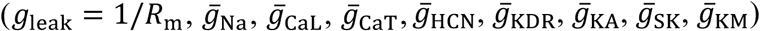, along with the decay time constant of calcium (***τ***_ca_; see below) were parameters that defined the intrinsic properties of the neuronal model. The reversal potentials for Na^+^, K^+^ and HCN channels were set at 55, –90 and –30 mV respectively.

The evolution of intracellular calcium as a function of calcium current (through voltage-gated calcium channels) and its buffering was modeled as in (Poirazi et al., 2003; Narayanan and Johnston, 2010; Honnuraiah and Narayanan, 2013):

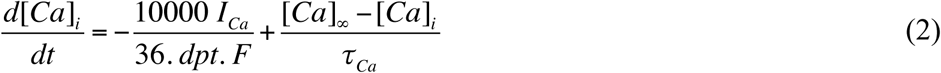

where *F* is Faraday’s constant, *I*_Ca_ is calcium current, the default value of the calcium decay time constant ***τ***_Ca_=30 ms, *dpt*=0.1 µm represented the depth of the shell, and [*Ca*]_∞_=100 nM defined the steady state value of cytosolic calcium concentration [*Ca*]_*i*_.

The current through the AMPA receptor (AMPAR) was modeled as the sum of currents carried by sodium and potassium ions (Narayanan and Johnston, 2010):

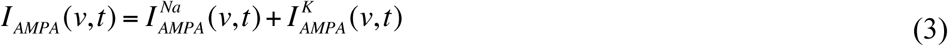

where,

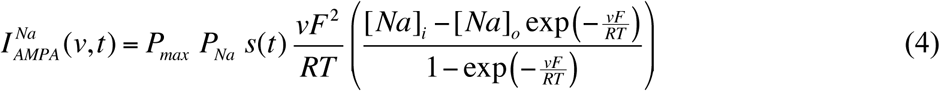

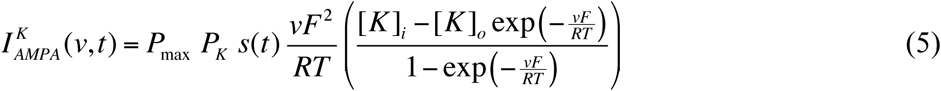

where *P*_max_ represented the maximum permeability of the AMPA receptor. The relative permeability ratios *P*_Na_ and *P*_K_ were equal and set to 1. *s*(*t*) was set as:

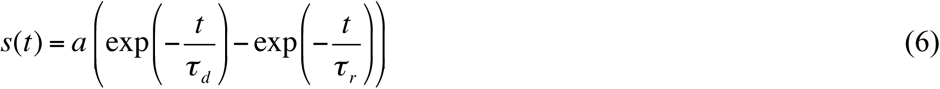

where *a* is a normalization constant, making sure that 0 ≤ *s*(*t*) ≤ 1, τ_r_, the parameter governing rise time was set to 2 ms and τ_d_, the decay time constant was 10 ms (Narayanan and Johnston, 2010). One hundred independently driven AMPAR synapses impinged on the neuron with independent presynaptic trains of action potentials stochastically activating these synapses based on an overall firing rate pattern (see below).

### Conductance-based synaptically driven inputs and population activity of place cells

Place-cell inputs were fed as probabilistic afferent activity impinging on the synapses described above. The frequency of place-cell inputs impinging as presynaptic afferents to these synapses was modeled as a Gaussian-modulated cosinusoidal distribution, with the frequency of the sinusoid set at 8 Hz. The presynaptic activation profile of our conductance-based synapses was derived from the simplified rate model, where place-cell inputs were modeled as Gaussian-modulated cosinusoidal *currents* (Geisler et al., 2010). This modification was essential because a current-based input would not account for the driving-force dependence of synaptic currents or the kinetics of receptors (Eq. 3–6). Therefore, the total afferent current was modeled to arrive through multiple conductance-based synapses whose presynaptic firing rates were stochastically driven. Specifically, with reference to the *n*^th^ place field within a linear arena, each synapse in a neuron received inputs with probability of occurrence at time *t* defined by (Fig. 1*C*):

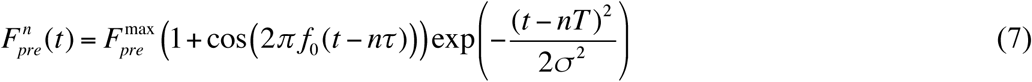

where *f*_0_ represented the cosine wave frequency (8 Hz) that translates to the intracellular theta frequency, 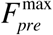 regulated the maximal input firing rate and σ defined the width of the Gaussian that controls the extent of the place field (Geisler et al., 2010). In this formulation (Fig. 1*E*), *T* signifies the longer time scale that corresponds to the temporal distance (the travel time) between adjacent place fields (modeled as a Gaussian) while τ characterizes a shorter theta time scale temporal difference between adjacent place fields (modeled as a phase shift in adjacent sinusoids at theta frequency). The standard deviation of the Gaussian distribution, σ, that governs the extent of single place fields was set as (Geisler et al., 2010):

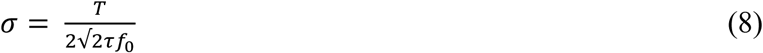

Within the TSC framework, the interference pattern between inputs from nearby place fields results in a reduction in the frequency of the extracellular theta or the population firing rate (Geisler et al., 2010). To construct the population firing rate (*f*_POP_) within our conductance-based model framework, we presented inputs from 50 distinct place field locations to synapses of a given model neuron (defined with specific intrinsic properties). Specifically, with reference Eq. 7 representing a linear arena traversal, *n* ∈ {1…50} reflects both a progressive shift in the center of the place field as well as a progressive phase shift in the theta time scale of individual place field inputs. The default values of *T* and τ were 180 ms and 10 ms, respectively.

For each value of *n* (∈ {1…50}), the synapses in the model neuron were stimulated stochastically with the stimulation probability of each synapse sampled from the distribution in Eq. (7). The firing patterns of the model neuron to each of these 50 place field traversals were computed (Fig. 1*D*). The spike times corresponding to each of these 50 place field traversals were derived from these firing patterns, and were converted to a binary time series (bin size 1 ms) indicating the presence or absence of a spike at a given time point. These binary time series were then summed across all 50 place field traversals to obtain the ensemble binary spike train that was then convolved with a Gaussian kernel to derive a smooth population firing rate profile (*f*_POP_; Fig. 1*F*). The Fourier transform of this population activity was computed and the peak in the Fourier magnitude spectrum (Fig. 1*F*, inset) was characterized as the population theta frequency (*f*_*θ*_), which represented the extracellular theta frequency (Geisler et al., 2010).

### Assignment of spike phases

In assigning spike phases with reference to the theta oscillation in the population firing rate (*f*_POP_), we first detected the troughs by determining the minima within each theta cycle (Fig. 1*F*). These detected troughs of the population theta were all assigned a phase of 0°. We used these detected troughs of the population theta to assign phase values to each spike corresponding to every place field input, with reference to the temporally aligned population theta oscillation. Specifically, let *t*_spike_ correspond to the timing of a spike corresponding to an arbitrary place field input. The population theta waveform was constructed from the ensemble output corresponding to all the 50 place field traversals, and encompasses the entire span of the linear arena (Fig. 1*F*). The spike patterns corresponding to each of these 50 distinct place field traversals, expectedly, span a much-restricted spatial (and temporal) extent, implying that each spike would have a temporally aligned stretch of the population firing rate waveform (Fig. 1*F–G*). Given this, for each *t*_spike_, we found two troughs of the population firing rate oscillation, one that immediately preceded *t*_spike_ (at time *t*_0_) another that immediately followed *t*_spike_ (at time *t*_1_). This implies that the neuronal spike (at *t*_spike_) occurred between two consecutive troughs (separated by a phase of 360°), at times *t*_0_ and *t*_1_, of the population firing waveform. Therefore the phase response *ϕ*_spike_ (in degrees) of the spike occurring at time *t*_spike_ with reference to *f*_POP_ was assigned as:

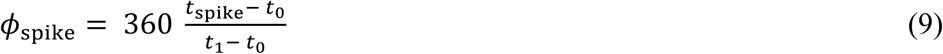

This assignment procedure was repeated for each spike corresponding to all the 50 place field traversals with temporally aligned *f*_POP_. Given the cyclical nature of phase precession, these phases were warped (by shifting phase values by a constant number) such that the representation has actual phase values around 360° at spatial locations closer to the beginning of the place field.

### Computation of Shannon’s entropy and mutual information

To define efficient phase coding, we took an information theoretic approach (Shannon, 1948) and used the mutual information between spatial stimuli and phase response to quantify efficiency. To do this, we extended the concept of efficient coding from sensory systems (Barlow, 1961; Bell and Sejnowski, 1997; Simoncelli and Olshausen, 2001; Lewicki, 2002; Simoncelli, 2003) and from single neurons (Stemmler and Koch, 1999) to phase coding within single place fields. Specifically, we chose maximization of mutual information to define an efficient phase code and explored the same from the perspective of efficient coding hypothesis, since it does not assume a parameterized functional form (Barlow, 1961; Bell and Sejnowski, 1995, 1997; Lewicki, 2002) for the phase code of the space. In our analyses, mutual information quantified the amount of information that the spike phase conveyed about the spatial stimulus that the neuron encountered, thereby signifying the ability of the model to clearly discriminate between various spatial stimuli. Mathematically, mutual information was defined as the difference between the response entropy and noise entropy (Shannon, 1948):

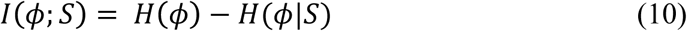

where *I*(*ϕ*; *S*) represented mutual information between stimulus (*S*; segregated into 20 distinct bins, where each bin constituted a different spatial stimulus) and neuronal phase response (*ϕ*) and *H*(*ϕ*| *S*) referred to the total noise entropy (Shannon, 1948). *H*(*ϕ*), the response entropy, was calculated as:

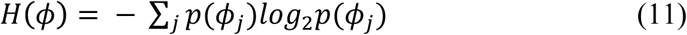

Where *p*(*ϕ*_*j*_), the probability of occurrence of the *j*^*th*^ phase bin (the 0–360° phase space was segregated into 360 bins; 0 ≤ *j* ≤ 359), was defined as:

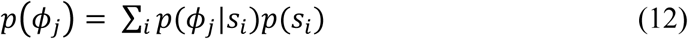

wherein the probability distribution of phases was derived by summing the conditional probability distributions of phases for various stimuli weighted by the probability of the stimulus. The total noise entropy was computed as:

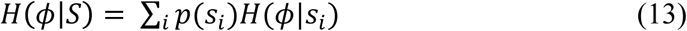

where *H*(*ϕ*|*S*_*i*_) represented the conditional noise entropy for stimulus *s*_*i*_, and was computed as:

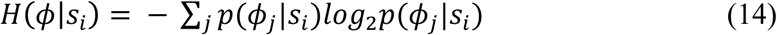

where *p*(*ϕ*_*j*_|*S*_*i*_) defined the conditional probability of *j*^*th*^ phase given *i*^*th*^ stimulus. Intuitively, response entropy captures the uncertainty in phase and noise entropy captures the uncertainty in phase despite the knowledge of the stimulus identity. Thus, noise entropy is that part of uncertainty that does not contribute any information about the stimulus and thereby is detrimental to information transfer. This explains mutual information as the difference between response and noise entropies (Shannon, 1948).

Efficiency in phase coding was analyzed on these theoretical grounds in order to understand the effectiveness of this temporal code in representing spatial information. In assessing the efficiency of spike phases corresponding to the 50 place field traversals (Eq. 9) obtained from our conductance-based model, we mapped the spike timings (and the corresponding spike phases) elicited for each place field traversal to a normalized place field space spanning 0 to 1. Specifically, with spike phase defined as the response and one-dimensional space constituting the stimulus, the construction of *p*(*ϕ*_*j*_|*S*_*i*_) (Eq. 14) requires phase responses to multiple spatial traversals. In our formulation, the 50 place field inputs correspond to distinct spatial traversals, and the corresponding spike phase responses are those of the same model to stochastic presynaptic inputs (Eq. 7) arriving onto the synapses. Spatially, these 50 place field traversals are different only in terms of the place field center shifting by *T* ms for every consecutive place field input, along with a ***τ*** ms phase shift on the theta scale (Eq. 7). Therefore, to construct *p*(*ϕ*_*j*_|*S*_*i*_), we superimposed the spike phase responses to multiple place field traversals by normalizing them with reference to their respective place field centers (Fig. 2).

**Figure 2.**
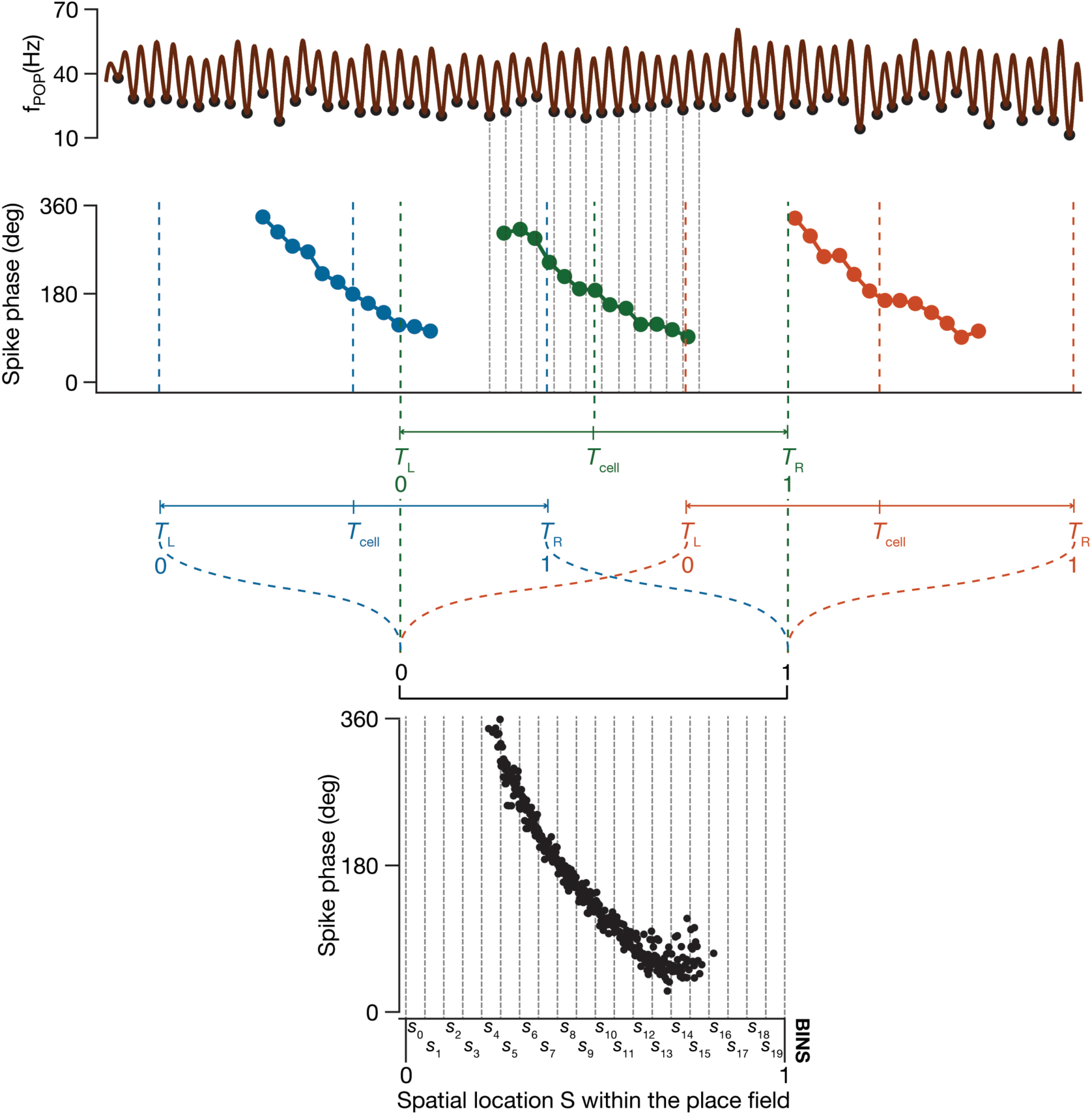
Illustration of computing the phase-space plot within our modeling framework. The population firing frequency (*f*_POP_) constituted the reference theta (first row, brown), which was computed from the spikes corresponding to all 50 place-field traversals. Spike phases (second row; blue, green and orange dots denote spike phases for three representative traversals) corresponding to each place field traversal were computed with reference to the detected troughs (top, black circles) of *f*_POP_. Each place-field traversal had a place field center (*T*_cell_), a left extreme (*T*_L_) and a right extreme (*T*_R_) beyond which there can be no spike corresponding to that traversal. *T*_cell_, *T*_L_ and *T*_R_ for each of the three representative place-field traversals are demarcated with color-matched dotted lines. The *T*_L_ to *T*_R_ extremes of each place field traversal are then mapped respectively to 0 and 1 on normalized spatial scale (*S*), with *T*_cell_ falling at 0.5 on this normalized scale. The spike phases corresponding to multiple traversals are then superimposed onto this normalized scale to yield the phase-space plot (third row). The spatial scale is split into 20 different bins (third row) for computing the probability distributions that are used to calculate the efficiency of spatial information transfer through this phase code.

Such normalization of spatial locations to 0–1 for each place field input, by accounting for their field center required an estimation of the spatial extent of each place field (so that no spike phases that belonged to the place field were omitted). The standard deviation (*σ*) of the place field Gaussian (Eq. 8) offered an ideal metric for determining this, and we employed *2σ* on either side of the respective place field center as the extent of each place field. With the center and extent of each place field known, the space normalization of each spike phase was defined by mapping the left and rights extremes of each place field to 0 and 1, respectively. Specifically, for the *n*^th^ (0 ≤ *n* ≤ 49) place field traversal, the field center (*T*_cell_) and the left (*T*_L_) and right (*T*_R_) extremes were (Fig. 2):

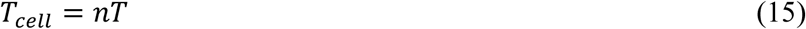

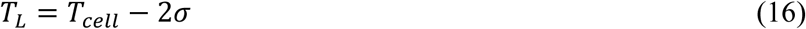

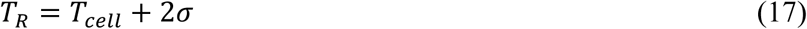

Therefore, the normalized spatial location (0 ≤ *S* ≤ 1) for a spike occurring at *t*_spike_ for a specific traversal was given as:

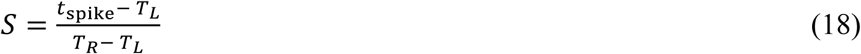

with *T*_L_ and *T*_R_ calculated for the specific traversal as in Eq. (15–17). As all spike phases (Eq. 9) corresponding to each of the place field traversals were now mapped onto a normalized spatial coordinate, all spike phases were superimposed to obtain a response (spike phase *ϕ*) *vs*. stimulus (normalized space *S*) plot which was employed for computing *p*(*ϕ*|*S*) (*e.g.*, Fig. 3*A–B*).

**Figure 3.**
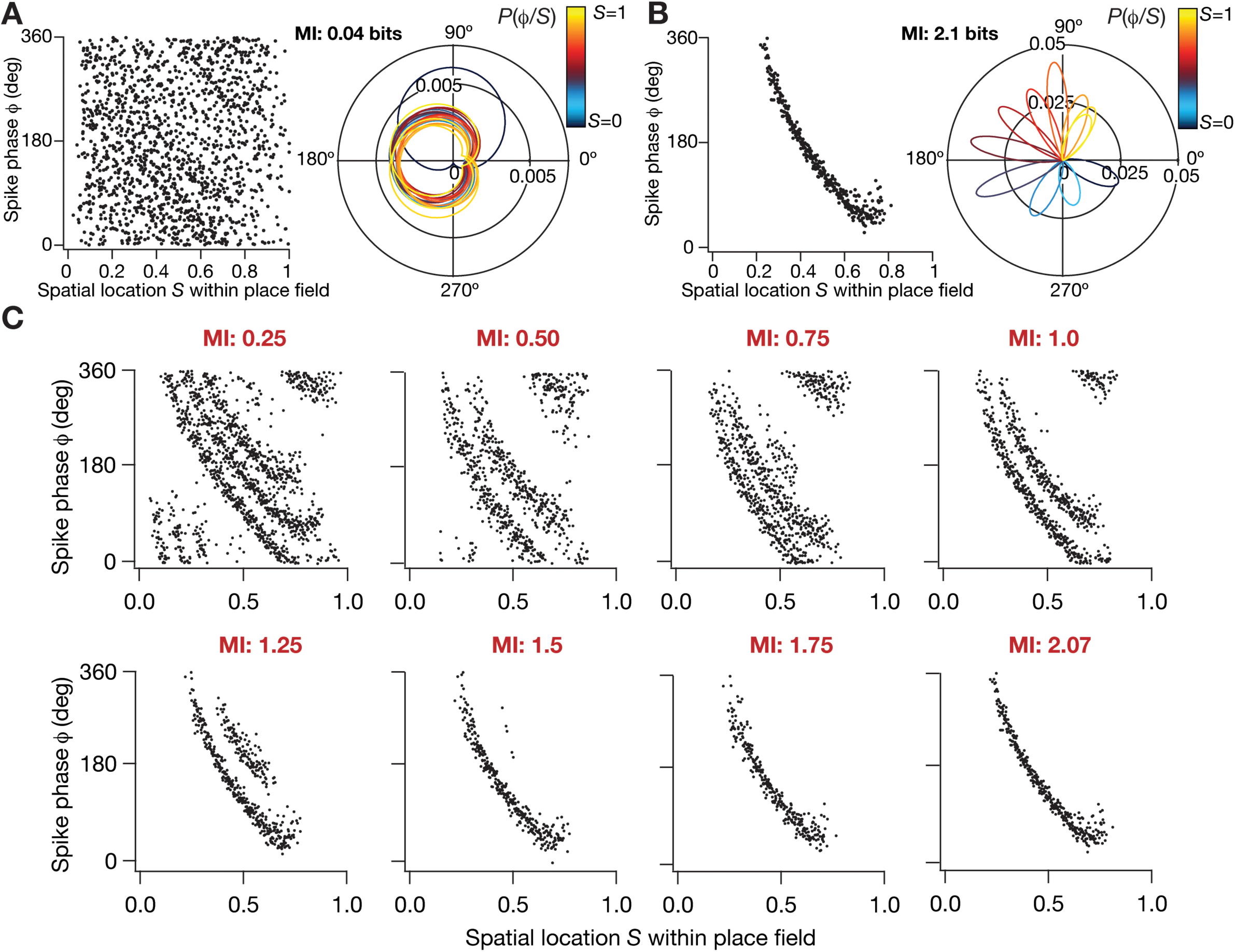
Spatial information transfer through spike phases was differential across models obtained from the unbiased stochastic search. (A–B) Changes in neuronal intrinsic properties are sufficient to alter phase precession and the efficiency of the phase code. Phase profile of a model neuron that failed to express phase precession (A, *Left*) despite receiving identical synaptic inputs, thereby lowering the efficiency of information transfer (mutual information between firing phase and spatial location=0.04 bits) through the phase code. A polar coordinate representation of conditional probabilities *P*(*ϕ*/*S*) of firing phases given the spatial bins within which the model received inputs (A, *Right*). Theta phase precession (B, *Left*) and conditional probability distribution of phase responses (B, *Right*) for another model neuron (B) with a higher efficiency of information transfer (mutual information between firing phase and spatial location=2.1 bits) through the phase code. The color-code correspond to the different spatial stimuli (bins), and the normalized color code is provided along with the *P*(*ϕ*/*S*) graphs. (C) Phase precession of eight example model neurons with successively higher mutual information values, picked from the pool of 11,000 models that were generated as part of the stochastic search procedure. The threshold for choosing efficient models was set at 1.5 bits of MI, implying that only the last three of the models above will be chosen as efficient models.

We computed *p*(*ϕ*|*S*) from this superimposed response-stimulus plot by binning the normalized stimulus axis *S* into 20 bins (Fig. 2), with each bin representing a spatial stimulus (*s*_i_, 0 ≤ *i* ≤ 19, in Eq. 12–14). In computing the conditional probability distribution *p*(*ϕ*|*s*_*i*_) (Eq. 12–14), we pooled all the spike phases that belonged to the bin *s*_i_ (bin *i* spanned *i*/20 to (*i*+1)/20 of the normalized stimulus axis *S*). We computed the mean and variance of the phases within each stimulus bin *s*_i_, and constructed a normal distribution with these statistics to yield *p*(*ϕ*|*s*_*i*_) (*e.g.*, Fig. 3*A–B*). Finally, the mutual information between stimulus (*S*) and neuronal phase response (*ϕ*), *I*(*ϕS*), was computed employing the conditional distribution *p*(*ϕ*|*s*_*i*_) and the probability of occurrence of each stimulus bin *p*(*s*_*i*_), which was considered to be a uniform distribution (implying uniform traversal of space), using equations 10–14.

In summary, the computation of *I*(*ϕ;S*) for a given conductance-based model neuron with a specified set of intrinsic properties (defined by parameters inherent to Eq. 1) entailed the computation of the spike phase responses for each of the 50 place field traversals (Eq. 7, Eq. 9), and computation of the response and noise entropies (Eq. 11–14) preceded by computation of *p*(*ϕ*|*S*) after normalization of the spatial coordinates (Eq. 15–18).

### Multiparametric multiobjective stochastic search: An unbiased global sensitivity analysis technique to explore parametric dependencies and degeneracy

Global sensitivity analysis is an algorithm employed to explore the entire parametric space and thereby alleviate the bias in interpretations that might result due to exploration across a narrow parametric regime. One methodology to perform global sensitivity analysis is to use a multiparametric multiobjective stochastic search (MPMOSS) technique that involves assigning a broad parametric space for every parameter in the model, followed by construction of individual models through uniform random sampling of all the parameters numerous times. As the choices of parameters are not biased, this constitutes an unbiased search strategy to assess the cross-dependencies of parameters, and to assess the emergence of degeneracy, the ability of disparate structural components to elicit similar functional outcomes. Models constructed through such stochastic search of a broad parametric space are then evaluated for their ability to match with multiple physiological constraints to assess their validity. Specifically, depending on the question in hand, the validation process involves the computation of model physiological measurements and matching them with corresponding experimental counterparts. Models that satisfy these physiological criteria are then classified as valid models and the underlying combination of parameters of all valid models are then employed for further analyses on parametric cross-dependencies and degeneracy (Foster et al., 1993; Prinz et al., 2004; Marder and Goaillard, 2006; Marder, 2011; Marder and Taylor, 2011; Rathour and Narayanan, 2012, 2014; Anirudhan and Narayanan, 2015; Mukunda and Narayanan, 2017; Basak and Narayanan, 2018; Mittal and Narayanan, 2018; Mishra and Narayanan, 2019).

To explore degeneracy in the emergence of efficient phase codes and to assess the role of intrinsic neuronal properties in the emergence of such codes, we performed MPMOSS on 11 parameters that are critical to our model. The 11 model parameters that we included in the stochastic search were the 9 maximal channel conductances 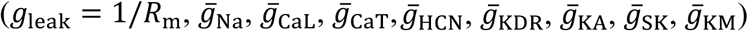,, the decay time constant of calcium (*τ*_Ca_) and synaptic permeability (*P*_max_). The exhaustive search space for each of these parameters is given in Table 1.

**Table 1.** Parameters and their ranges governing the multiparametric multiobjective stochastic search.

| Parameter | Minimum value | Maximum value |
| --- | --- | --- |
| Sodium conductance, $\bar{g}_{Na}$ (mS/cm<sup>2</sup>) | 1 | 100 |
| Delayed rectifier potassium conductance, $\bar{g}_{KDR}$ (mS/cm<sup>2</sup>) | 0.1 | 100 |
| A type K<sup>+</sup> conductance, $\bar{g}_{KA}$ (mS/cm<sup>2</sup>) | 0 | 10 |
| L type Ca<sup>2+</sup> conductance, $\bar{g}_{CaL}$ (μS/cm<sup>2</sup>) | 0 | 100 |
| Calcium gated k+ conductance, $\bar{g}_{SK}$ (μS/cm<sup>2</sup>) | 0 | 100 |
| HCN conductance, $\bar{g}_h$ (μS/cm<sup>2</sup>) | 0 | 100 |
| M type K<sup>+</sup> conductance, $\bar{g}_{KM}$ (μS/cm<sup>2</sup>) | 0 | 100 |
| T type Ca<sup>2+</sup> conductance, $\bar{g}_{CaT}$ (μS/cm<sup>2</sup>) | 0 | 100 |
| Calcium decay constant, $\tau_{Ca}$ (ms) | 20 | 100 |
| Specific membrane resistance, $R_m$ (kΩcm<sup>2</sup>) | 20 | 60 |
| Synaptic permeability, $P_{max}$ (nm/s) | 0.2 | 2 |

Each of the 11 parameters in this multiparametric space (Table 1) was uniformly sampled 11,000 times to generate as many unique models, and *I*(*ϕ;S*) was computed for each of these 11,000 conductance-based models (each with distinct intrinsic properties) employing the procedure outlined in the previous section. As our goal was to elucidate information efficiency in the phase code, the first validation criterion that we imposed on these models was based on *I*(*ϕ;S*). Specifically, all models that had a high value (>1.5 bits; Fig. 3*C*) of this mutual information were considered to be “MI-valid” models. Of the 11000 models that were constructed, 284 models were classified as MI-valid models (∼2.6%). The distribution of parameters in these valid models was then analyzed to assess degeneracy, parametric cross-dependencies and the role of intrinsic neuronal properties on information efficiency of the phase code.

### Global sensitivity analysis to explore degeneracy in the concomitant emergence of efficient phase coding and robust excitability measurements

Whereas mutual information based validation identified efficient models, this validation procedure did not account for whether the model was endowed with excitability measurements akin to CA1 pyramidal neurons. To do this, we imposed a second layer of validation criteria on the 284 MI-valid models by computing their physiological measurements of intrinsic excitability and validating them against their electrophysiological counterparts. These intrinsic measurements were resting membrane potential (RMP), standard deviation of RMP (σ_RMP_) to avoid fluctuations (consequent to channel interactions) in the emergence of a resting state, input resistance (*R*_in_), firing rate at 50 pA (*f*_50_) and firing rate at 250 pA (*f*_250_).

To measure RMP, the model neuron was allowed to achieve steady state (without injection of any current and without activation of any of the afferent synapses) for a period of 6 seconds (Fig. 5*B*). The distribution of membrane potential values over the last 1-second (*i.e.*, 5–6 s period) was employed to compute its mean and standard deviation, which were then defined as RMP and σ_RMP_ respectively. We injected the model neuron with pulse currents (each for 500 ms after steady-state RMP was achieved as above) of amplitudes ranging from –50 pA to 50 pA (with incremental steps of 10 pA each) and the resulting steady-state voltage response was recorded for each amplitude of current (Fig. 5*C*). The steady-state voltage deflection from RMP was plotted as a function of injected current amplitude. The slope of the linear fit to this *V*–*I* plot was defined as the input resistance of the model neuron (*R*_in_). Finally, the firing rates at 50 pA and 250 pA were measured by injecting constant pulse currents of the respective magnitudes and computed as the number of action potentials elicited during a 1-second period. All these intrinsic measurements were made after allowing the RMP to stabilize for 6 seconds. These intrinsic measurements were computed for each of the 284 models that were then validated based on experimentally derived ranges of measurements (Table 2). This process left us with 132 models that were both MI-valid and intrinsically valid. The distribution of parameters in these valid models was then analyzed to assess degeneracy, parametric cross-dependencies and the role of intrinsic neuronal properties in the concomitant emergence of efficient phase codes and robust intrinsic excitability.

**Table 2.** Electrophysiological bounds on intrinsic measurements employed for model validation.

| **Intrinsic measurement** | **Lower bound** | **Upper bound** |
| --- | --- | --- |
| Mean Resting membrane potential, $\mu_{\text{RMP}}$ (mV) | -75 | -60 |
| Standard deviation of resting membrane potential, $\sigma_{\text{RMP}}$ (mV) | < 1 | |
| Input resistance, $R_{\text{in}}$ (MΩ) | 30 | 120 |
| Firing rate at 50 pA, $f_{50}$ (Hz) | = 0 | |
| Firing rate at 250 pA, $f_{250}$ (Hz) | 1 | 45 |

**Figure 4.**
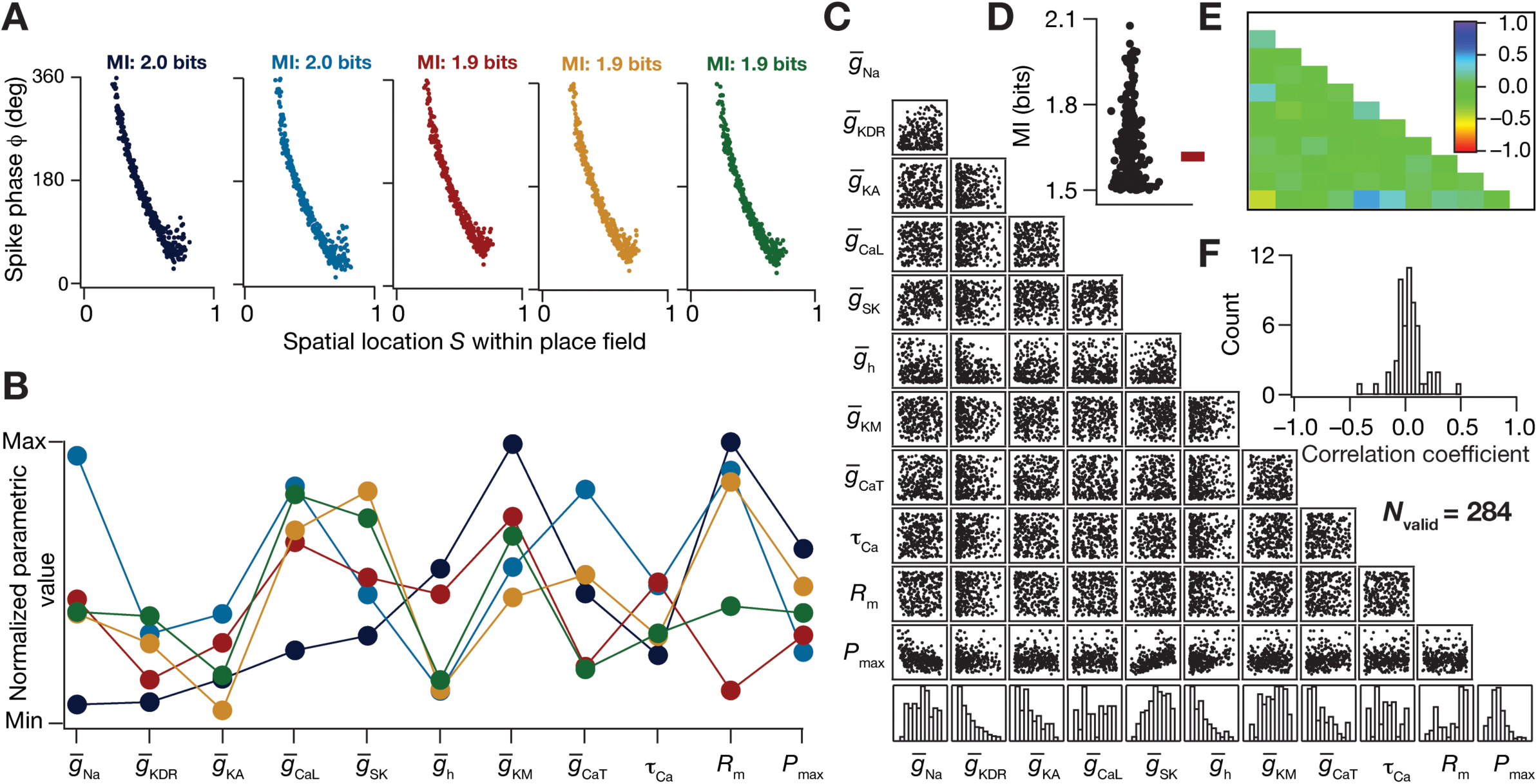
Degeneracy in efficient phase coding achieved through disparate neuronal intrinsic properties. (A) Phase precession of five example model neurons with very similar mutual information values, picked from a pool of 284 models classified as efficient phase coders due to an MI value greater than 1.5 bits. (B) Distribution of active and passive parameters that defined the five example models normalized between their respective ranges. (C) Scatter plot matrix of all the 11 parameters that govern the 284 valid models, displaying pairwise correlations. The last row shows the histogram of the 11 parameters that defined these model neurons. (D) Beeswarm plot of the mutual information between firing phase *ϕ* and spatial location *S* for all the 284 valid models. (E) Pearson’s correlation coefficient quantifying the pairwise correlations, of scatter plots shown in panel (C). (F) Histogram of pearson’s correlation coefficients of all the 11 parameters, clustering around zero.

**Figure 5.**
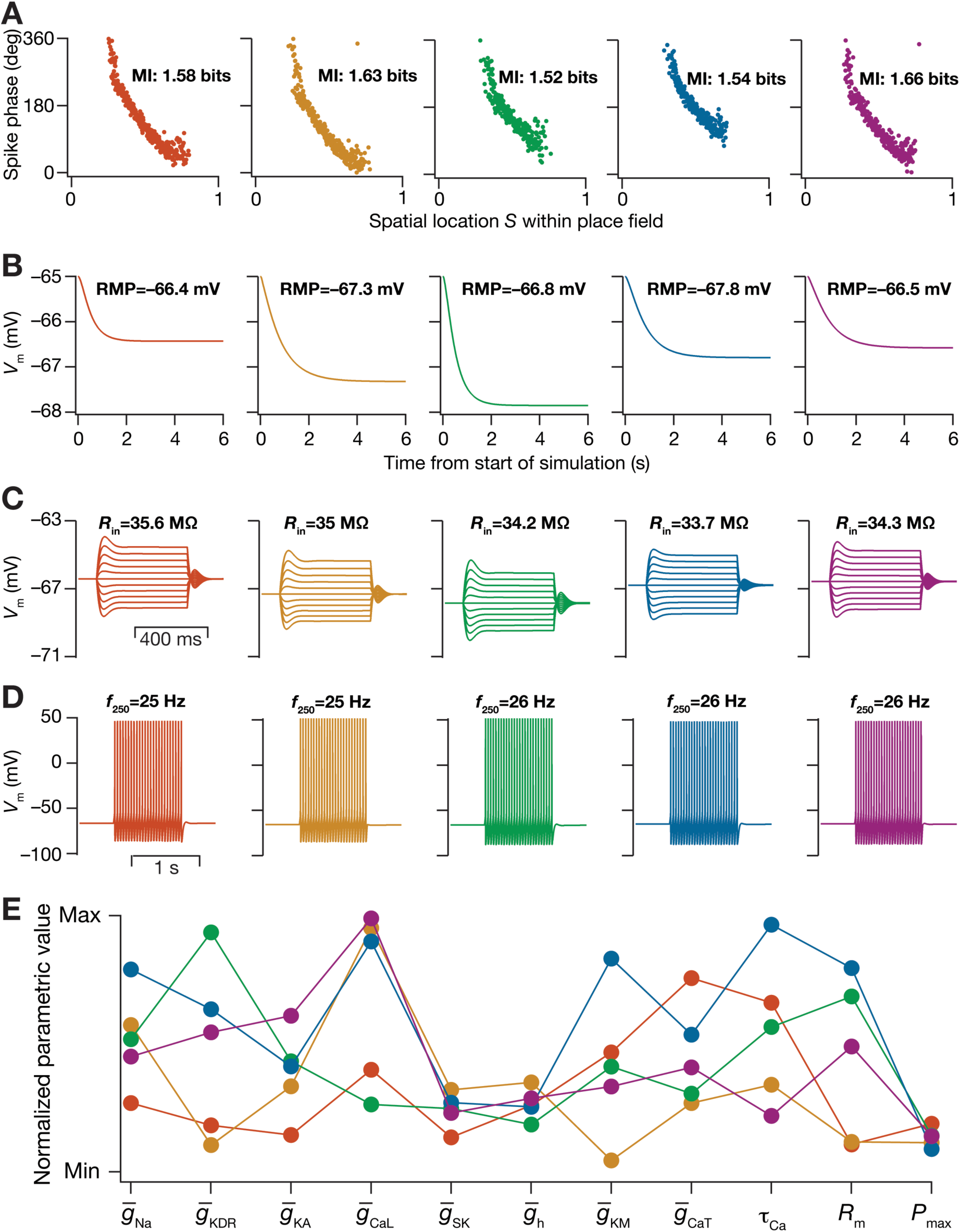
Disparate parametric combinations resulted in the concomitant emergence of efficient phase coding and excitability homeostasis. (A–D) The five color-coded columns represent different model neurons picked from a pool of 132 models that were classified as valid, based on high MI values and their intrinsic measurements satisfying electrophysiological bounds specified in Table 2. The five models were chosen based on similarity in MI values as well as intrinsic measurements. The similarity of phase precession curves and mutual information (A), resting membrane potential (B), input resistance (C) and firing rate at 250 pA (D) across these five models may be noted. (E) Normalized parameter values that yielded the five models represented in (A–D), color-coded to represent the model identity. The parameter values are seen to span a large range (E), despite similarities spanning efficient encoding (MI) and intrinsic (RMP, *R*_in_, *f*_250_) measurements. The firing rate for 50 pA current injection, *f*_50_, was identically zero and *σ*_RMP_ < 0.01 mV for all 132 valid models (Table 2).

**Figure 6.**
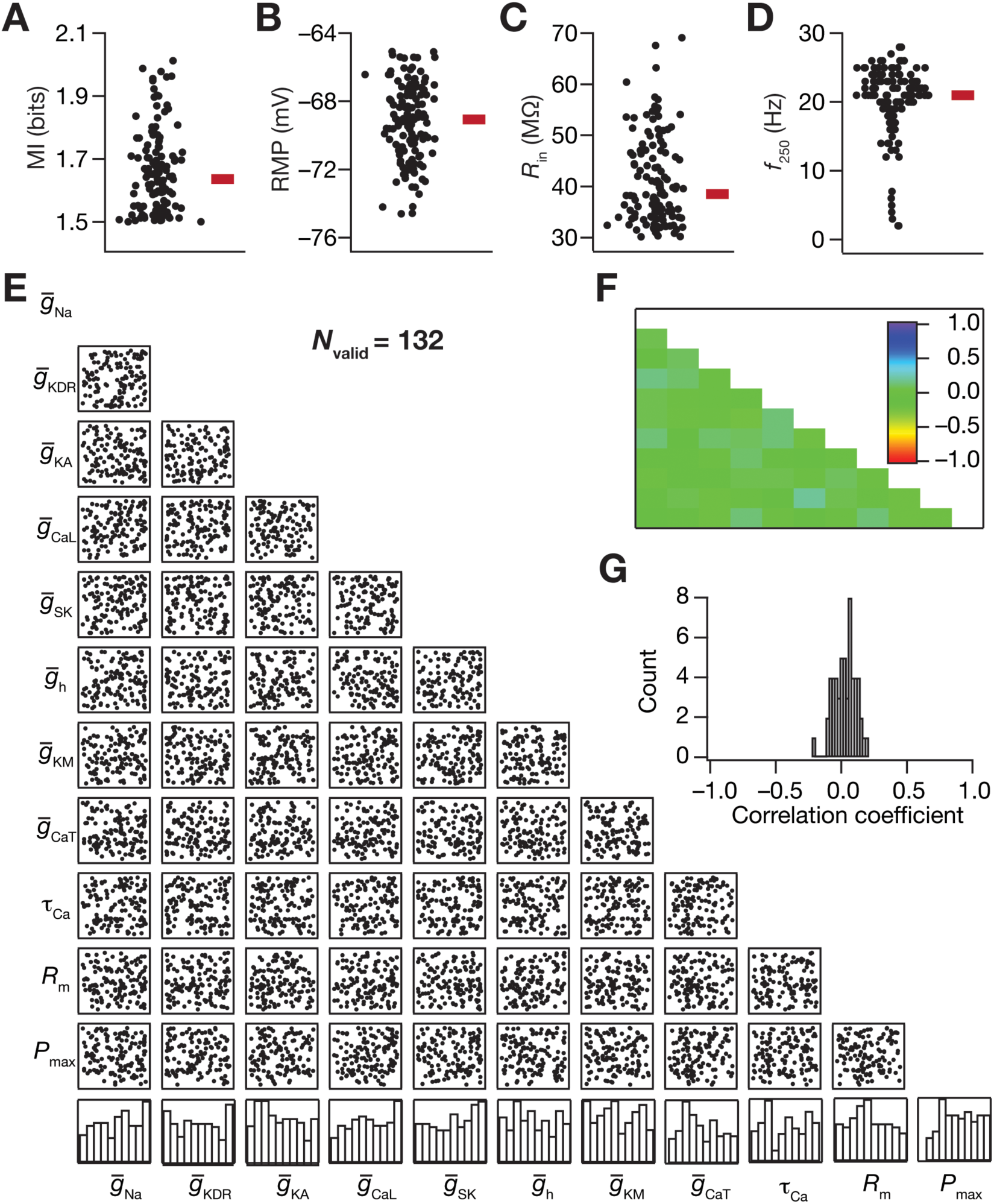
Synergistic interactions between synaptic and intrinsic properties drive the concomitant emergence of efficient phase coding and excitability homeostasis within the framework of degeneracy. (A–D) Beeswarm plots of the mutual information between firing phase *ϕ* and spatial location *S* (A), resting membrane potential (B), input resistance (C) and firing rate for 250 pA (*f*_250_) current injection (D) for all the 132 valid models. (E) Scatter plot matrix showing pairwise correlations between parameters that underlie 132 models that were classified as valid based on MI values and excitability measurements. (F) The Pearson correlation coefficient matrix for all the pairwise correlations in the scatter plot matrix in (E). (G) Histogram of correlation coefficients shown in (F).

**Figure 7.**
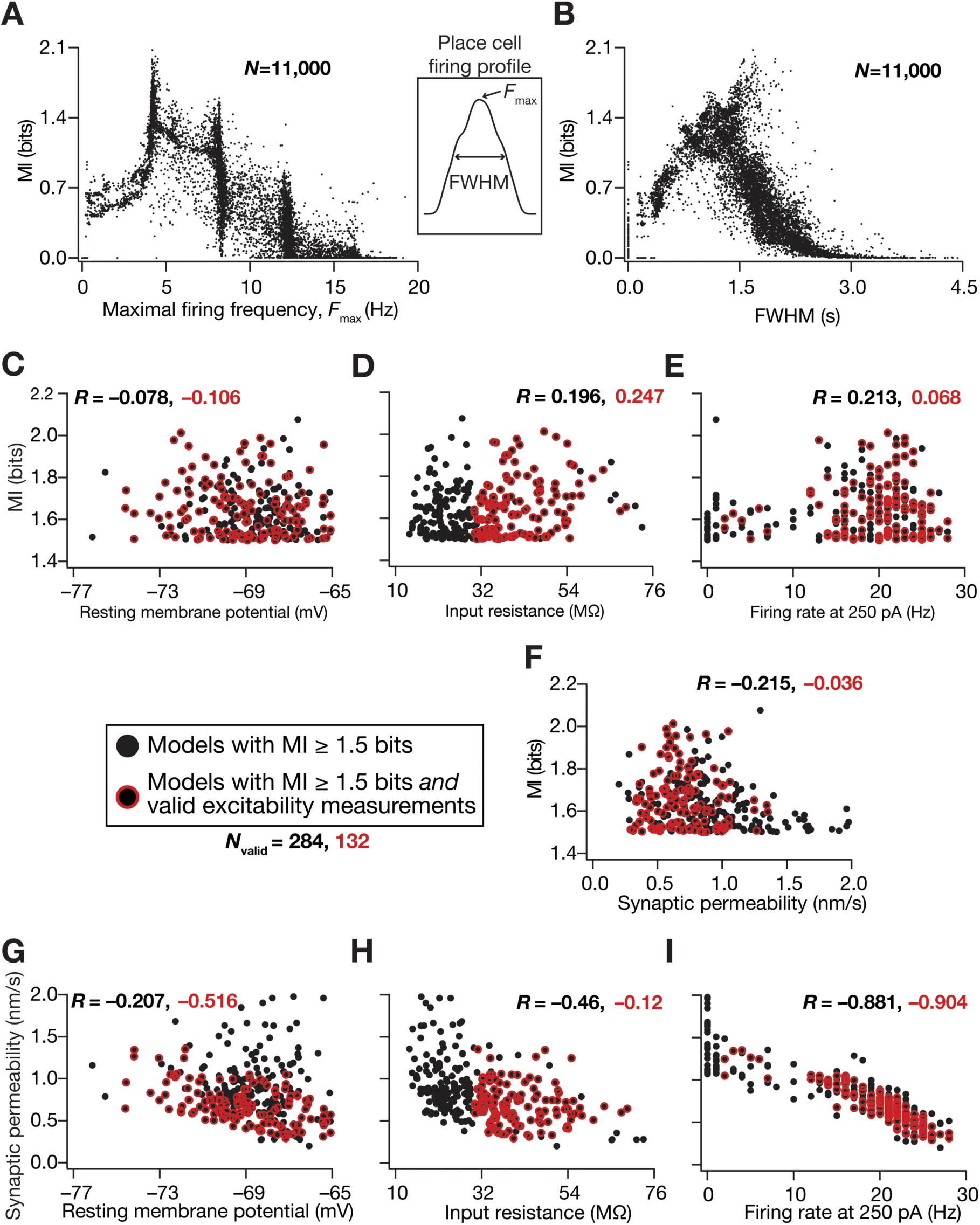
Synergistic functional interactions between synaptic strength and intrinsic excitability governed the emergence of efficient phase coding. (A–B) Mutual information plotted as a function of the respective maximal firing rate (*F*_max_) and full-width at half maximum (FWHM) of all 11,000 models. Inset between (A) and (B) shows a firing rate profile indicating *F*_max_ and FWHM of an example place-cell firing rate profile. (C–F) Scatter plots of mutual information *vs*. resting membrane potential (C), input resistance (D), firing rate at 250 pA (E) and synaptic permeability (F) unveiled the absence of strong correlations between efficient coding and intrinsic/synaptic functional properties. (G–I) Scatter plots of synaptic permeability *vs*. resting membrane potential (G), input resistance (H) and firing rate at 250 pA (I) revealed strong correlations between intrinsic firing frequency and synaptic strength (I). For all panels, black circles represented the population of models (*N*_valid_=284; from Fig. 3) that were endowed with an MI ≥ 1.5 bits, and a red outline around the black circles represented a subset of these models that were also endowed with valid intrinsic measurements (*N*_valid_=132; from Fig. 6). The values against *R* represent Pearson’s correlation coefficients for the respective color-coded scatter plots.

**Figure 8.**
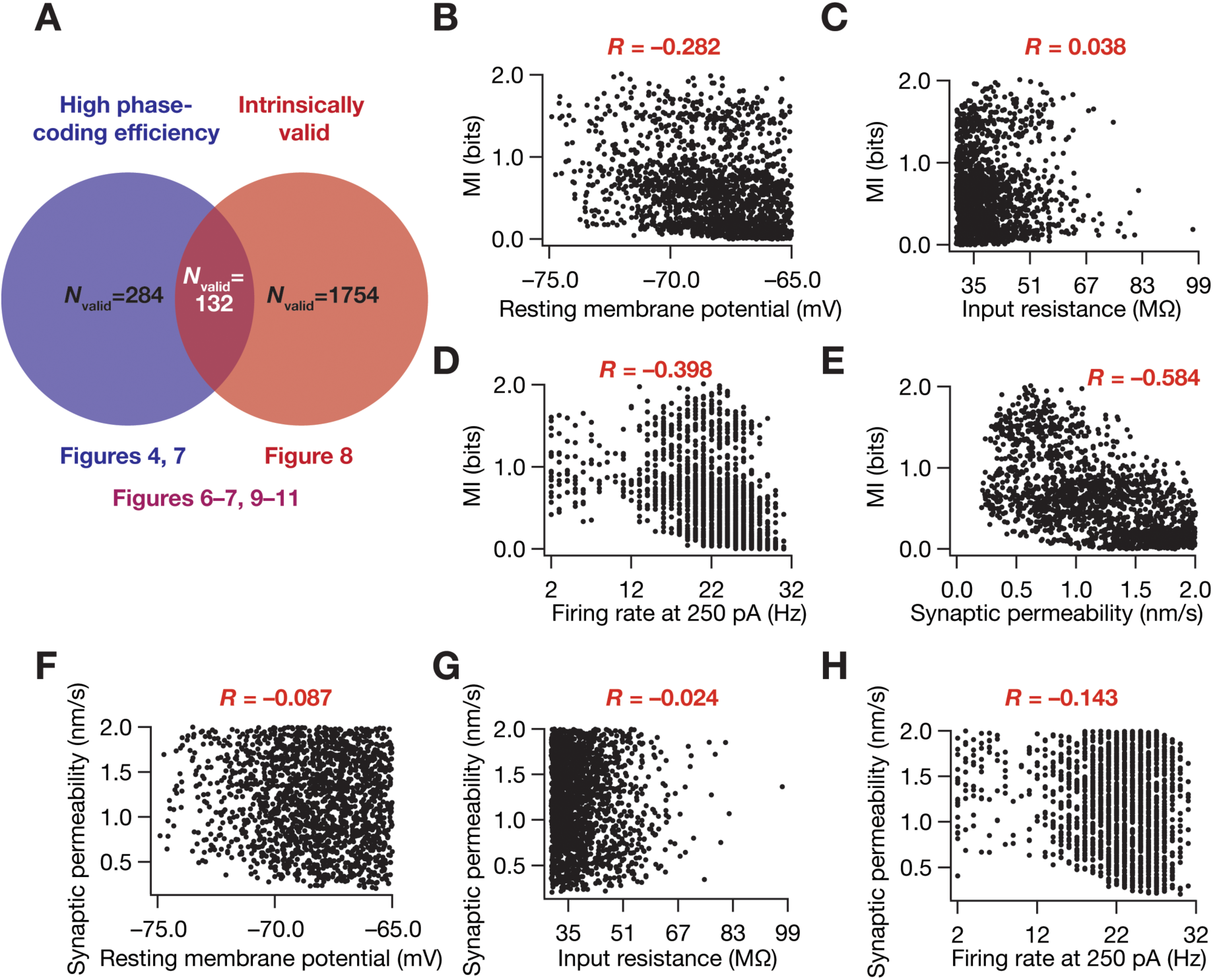
Efficiency in phase coding was not strongly correlated with intrinsic excitability or synaptic strength in models with signature excitability characteristics of CA1 pyramidal neurons. (A) For other analyses in the study, as a first step, we had found efficient models with high MI (*N*_valid_=284; blue circle) from the 11,000 stochastically generated models (Fig. 3). As a second step in such analyses, we found models with concomitant signature excitability characteristics as a *subset* (*N*_valid_=132; purple intersection region between the red and blue circles) of these high-MI efficient models (Figs. 6–7). For the analyses presented in this figure, instead of finding efficient models as the first step, we found models among the entire set of 11,000 stochastically generated models that satisfied signature excitability characteristics. This validation procedure yielded 1754 models that were intrinsically valid (red circle), but did not have any constraints imposed on their efficiency (mutual information). The analyses presented in this figure correspond to these 1754 intrinsically valid models. (B–E) Scatter plots of mutual information *vs*. resting membrane potential (B), input resistance (C), firing rate at 250 pA (D) and synaptic permeability (E) unveiled the absence of strong correlations between efficient coding and intrinsic/synaptic functional properties. (F–H) Scatter plots of synaptic permeability *vs*. resting membrane potential (F), input resistance (G) and firing rate at 250 pA (H) revealed the absence of strong correlations between intrinsic properties and synaptic strength. The values against *R* represent Pearson’s correlation coefficients for the respective color-coded scatter plots. *N*_valid_=1,754 for all plots.

**Figure 9.**
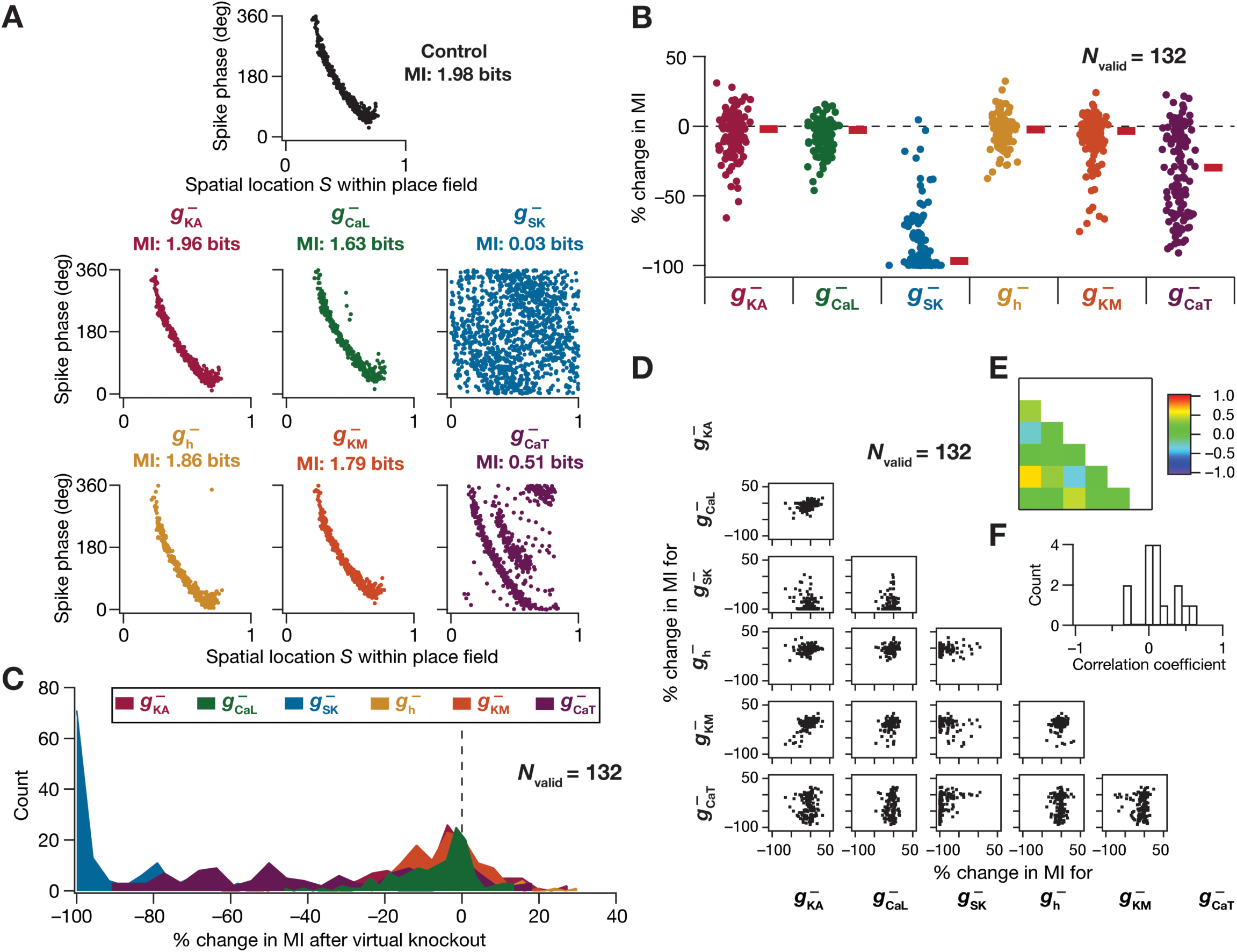
Virtual knockout models unveil differential and variable dependence of efficient phase coding on individual ion channels. (A) Phase precession profiles of an example model randomly picked from the 132 models that exhibited efficient phase coding and excitability homeostasis. Phase precession of this model with all the conductances intact (marked “Control”) and after virtual knockout of each of the 6 channels one at a time (the specific channel knocked out is mentioned on top of the respective phase-space plots). (B–C) Beeswarm plots (B) and histograms (C) of percentage changes in mutual information values after virtual knockout of each channel from the population of 132 models. (D) Scatter plot matrix showing pairwise correlations between percentage changes in MI after virtual knockout of the six distinct channel subtypes in all 132 models. (E) The Pearson correlation coefficient matrix for all the pairwise correlations in the scatter plot matrix in (D). (F) Histogram of correlation coefficients shown in (E).

**Figure 10.**
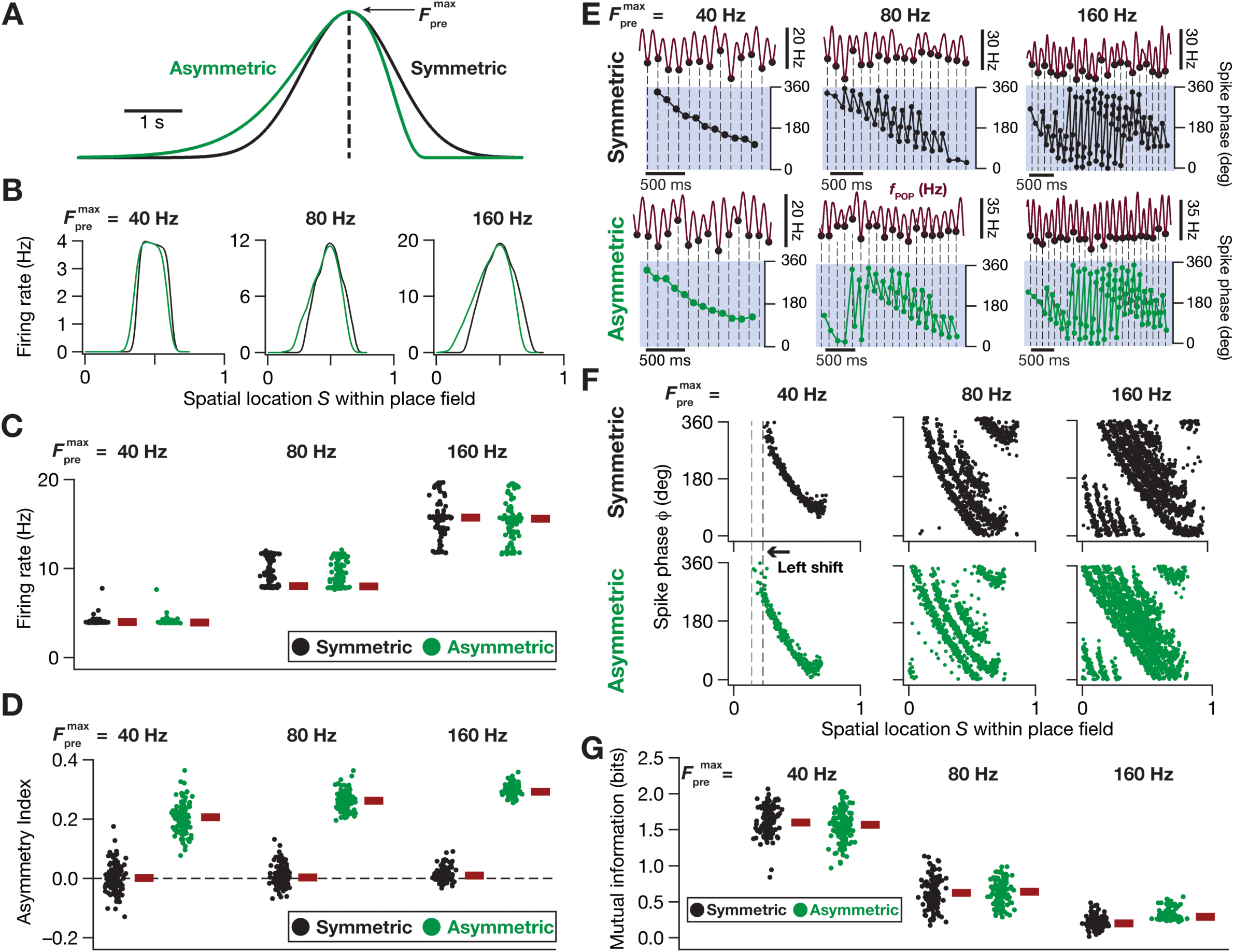
Experience-dependence of rate and phase coding modeled through an asymmetric ramp input. (A) Symmetric (black) and Asymmetric (green) profiles defining the probability distribution governing the activation of synaptic inputs arriving onto a model neuron during place field traversals. The symmetric profile is a theta-modulated Gaussian distribution while the asymmetric profile is a theta-modulated Erlang distribution. The peak, 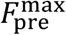, denotes the maximal pre-synaptic firing rate and the dashed black line indicates the center of the place field. (B) The three columns represent the firing rate profiles of an example model neuron that received symmetric (black) and asymmetric (green) inputs within its place field at three different pre-synaptic firing rates (40, 80 and 160 Hz). The place field extent is normalized between 0 and 1. (C–D) Beeswarm plots of firing rates (C) and asymmetry indices (D) of all the 132 valid models receiving both symmetric (black) and asymmetric (green) input profiles at three different pre-synaptic firing rates (40, 80, 160 Hz). (E) Single-trial phase precession plots of the same model neuron as shown in (B) with reference to the theta oscillation. The three columns represent the model neuron receiving three different pre-synaptic firing rates (40, 80 and 160 Hz) and the two rows indicate symmetric (black) and asymmetric (green) cases of synaptic inputs. The black filled dots (in top traces) and dashed lines represent the troughs, which are aligned in time with the spikes of the model neuron (dots, bottom traces), in each panel. Note that the temporal scale bars for 500 ms become progressively shorter with increase in 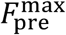. (F) Multi-trial phase precession plots of the same model neuron as shown in B) and (E) for symmetric (black, top) and asymmetric (green, bottom) cases at three different pre-synaptic firing rates (40, 80 and 160 Hz) shown in the three columns. The leftward shift of the phase code for an asymmetric synaptic input profile may be noted for all three different pre-synaptic firing rates. (G) Beeswarm plots of mutual information between spike phase and spatial location within the place field, for all the 132 valid models, for both symmetric (black) and asymmetric (green) cases at three different pre-synaptic firing rates (40, 80 and 160 Hz).

### Virtual knockout models

Within the MPMOSS framework, virtual knockout analysis is a technique that helps understand the sensitivity of the disparate valid models towards a specific ion channel conductance (Rathour and Narayanan, 2014; Anirudhan and Narayanan, 2015; Basak and Narayanan, 2018; Mittal and Narayanan, 2018). Virtual knockout analyses were performed by comparing model outcome before and after setting each of the 6 channel conductances 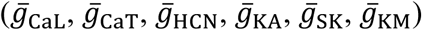 to zero. These simulations were performed on the 132 concomitantly MI- and intrinsically-valid models by knocking out the 6 channel conductances, one at a time. For each of the 132 models, and for each of the 6 channel conductances, we computed the phase code and associated *I*(*ϕ;S*) (132 × 6 = 792 simulations involving Eq. 7–18) to compare the impact of each channel on the efficiency of the phase code.

### Experience dependent asymmetry

To study experience-dependence of rate and phase codes, we used an asymmetric probability distribution that governed the activation of 100 independent AMPA synapses. Specifically, the probability of activation of afferent synapses was earlier defined by a symmetric Gaussian modulated cosinusoidal function (Eq. 7). To mimic experience-dependent asymmetry in the afferent ramp (Mehta et al., 1997; Mehta et al., 2000; Harvey et al., 2009), we replaced the symmetric Gaussian envelope by a horizontally-reflected Erlang distribution to construct the asymmetric envelope (Fig 10*A*). With this formulation, each synapse in the model neuron received inputs whose probability of occurrence, as a function of time was defined by an Erlang-modulated cosinusoidal distribution:

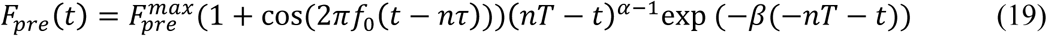

where all common parameters with Eq. 7 were identical in their description and function, and parameters α (=4) and β (=0.002) governed the extent of asymmetry.

In comparing the implications of symmetric and asymmetric afferent activations on the place-field rate and phase codes, we first constructed the firing rate profiles across each of the 50 different symmetric (Eq. 7) or asymmetric (Eq. 19) place field traversals for a given model. To do this, the spike time responses of each model neuron to the fifty distinct symmetric or asymmetric place field inputs were convolved with a Gaussian kernel to produce instantaneous firing rate responses. These fifty firing rate profiles were finally averaged to produce the rate code of that particular model neuron (*e.g.*, Fig 10*B*), in response to either the symmetric or the asymmetric input profiles. To assess the sensitivity of the model to afferent input strength, we presented either symmetric or asymmetric inputs at three different maximal pre-synaptic firing rates (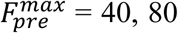 or 160 Hz). We computed an asymmetry index (*AI*) to assess the extent of asymmetry of the firing rate profile of a given model neuron:

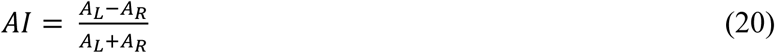

where *A*_L_ denoted, for each model neuron, the area under the firing rate profile to the left of the place field center (0 to 0.5 on the normalized spatial axis) while *A*_R_ represented the same to the right of the place field center (0.5 to 1 on the normalized spatial axis). The firing rate profiles and their asymmetry indices were computed for each of the 132 concomitantly MI- and intrinsically-valid models, at three different pre-synaptic firing rates, for both symmetric and asymmetric input profiles (Fig. 10*C–D*).

For computing the phase codes and associated *I*(*ϕ;S*) for a given model, with reference to 50 consecutive asymmetric place field inputs, we employed the procedure outlined earlier for symmetric inputs (Eq. 7–18), except with Eq. 7 being replaced by Eq. 19. We performed the computation of phase codes and associated *I*(*ϕ;S*) for symmetric or asymmetric inputs at three different maximal pre-synaptic firing rates (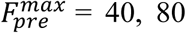 or 160 Hz). We compared phase codes and associated efficiency for all the 132 concomitantly MI-and intrinsically-valid models, with each model receiving either symmetric *vs*. asymmetric place field inputs.

### Analyses of theta power and ramp in neuronal voltage responses recorded during place field traversal

We employed the following measures for quantifying theta modulation and sub-threshold ramp in neuronal voltage responses during place-field traversal (Fig. 11): (a) area under the curve (AUC) of the Fourier spectrum of the theta-filtered signal; (b) subthreshold membrane potential ramp amplitude and (c) asymmetry index of the sub-threshold voltage ramp. Theta modulation in the voltage response was assessed by band-pass filtering the raw membrane potential signal between 4–10 Hz with a Butterworth filter of order 10. To quantify theta power, we computed the area under the Fourier spectrum of this filtered signal. To compute sub-threshold ramp, a place-cell signature at a subthreshold level (Harvey et al., 2009), we median filtered the raw voltage trace with a window of 1.75 s to derive the subthreshold voltage ramp. We computed the amplitude of the sub-threshold ramp as the difference between the peak and baseline voltage values of this filtered signal (Basak and Narayanan, 2018). We computed the asymmetry index (*AI*) to quantify the extent of asymmetry of the subthreshold voltage ramp exhibited by the model neurons as:

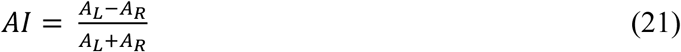

where *A*_L_ denoted the area under the median filtered voltage trace to the left of the place field center (which was computed for the chosen place field traversal *i*, as *T*i*) while *A*_R_ denoted the same to the right of the place field center.

**Figure 11.**
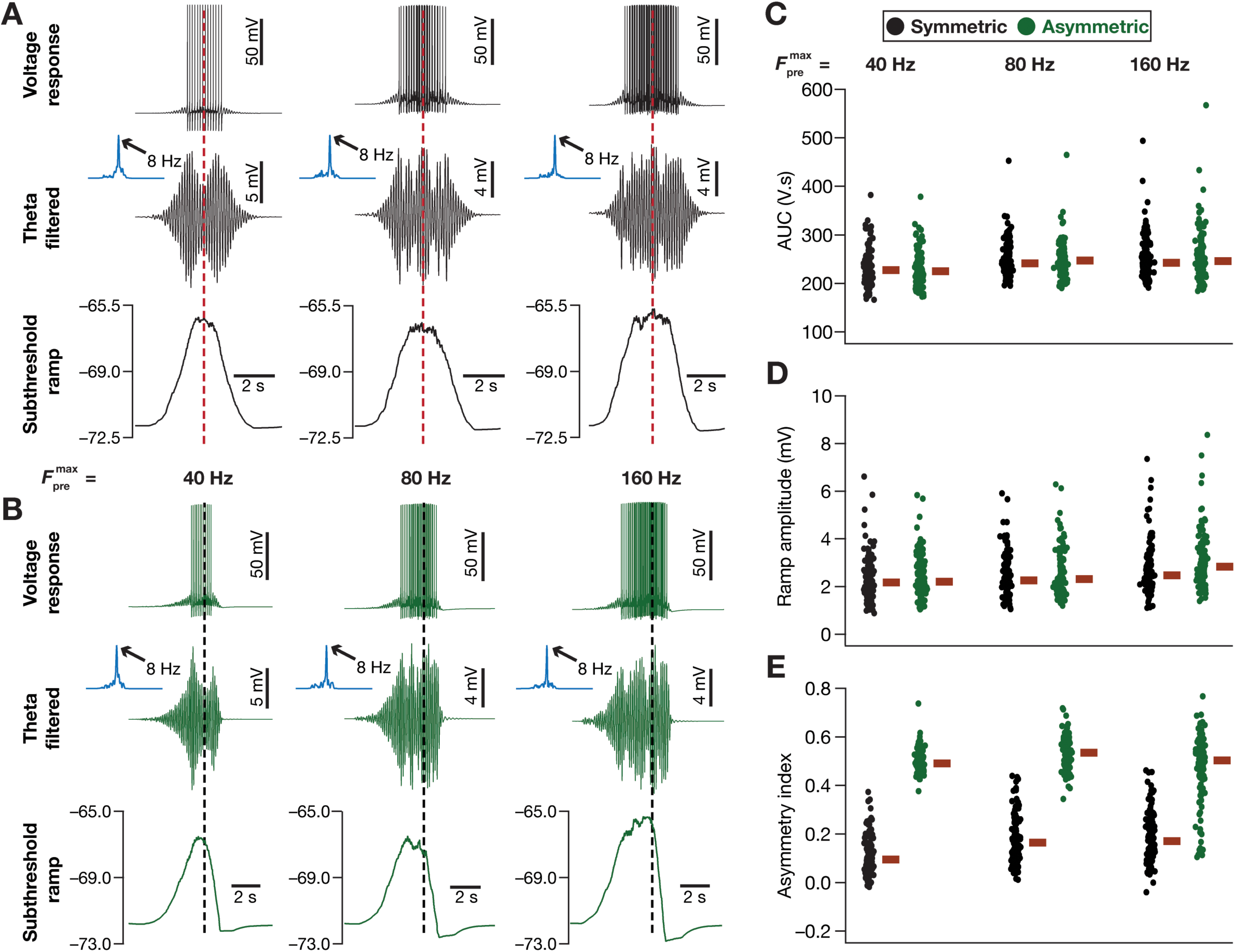
Theta modulation and subthreshold ramp exhibited by the model. For all panels in this figure, black and green correspond to symmetric and asymmetric synaptic activation profiles, respectively. (A–B) The three columns for all plots represent three different values of the maximal firing rate of the presynaptic input, 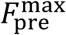 (40, 80 and 160 Hz from left to right). *Top*, Unfiltered voltage response of the model neuron for the symmetric (A) and asymmetric (B) synaptic activation during a place-field traversal. *Middle*, The band-pass (4–10 Hz, theta-band) filtered voltage response, represented in time domain along with its Fourier spectrum (blue) presented as an inset for symmetric (A) and asymmetric (B) cases. The area under the curve (AUC) for the blue Fourier spectrum traces in (A) were computed to be 234, 281 and 303 V.s, and for those in (B) were computed to be 253, 290 and 287 V.s, respectively for 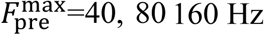. *Bottom*, The sub-threshold voltage response profile was obtained by median filtering the voltage response, for the symmetric (A) and asymmetric (B) cases. The dashed red lines in panel (A) and the dashed black lines in panel (B) represent the respective centers of the place fields. The traces shown in A–B are from the same model cell employed in Fig. 10*B*. (C–E) Beeswarm plots of AUC of the Fourier spectrum of the band-pass filtered signal (C), the sub-threshold ramp amplitude (D) and asymmetry index of the sub-threshold ramp (E) for all the 132 models. Measurements corresponding to symmetric (black) and asymmetric (green) synaptic activation profiles for the three values of 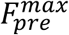 (40, 80 and 160 Hz, left to right) are shown. The red rectangles represent the corresponding median values.

### Computational details

All simulations were done using the NEURON programming environment (Carnevale and Hines, 2006), at 34°C with the simulation step size set at 25 µs. Data analyses and graph plotting were performed using Matlab and custom written software in the IGOR Pro environment.

## RESULTS

### Development of conductance-based model for phase coding within the temporal sequence compression framework

We developed a conductance-based model for phase precession within the TSC framework (Skaggs et al., 1996; Dragoi and Buzsaki, 2006; Geisler et al., 2010), building upon the rate-based model presented by Geisler and colleagues (Geisler et al., 2010). Our model was endowed with 8 different active ion channels (Fig. 1*B*) whose kinetic properties were derived from electrophysiological recordings from CA1 pyramidal neurons (Magee and Johnston, 1995; Sah and Isaacson, 1995; Hoffman et al., 1997; Magee, 1998; Migliore et al., 1999; Sah and Clements, 1999; Poolos et al., 2002; Migliore et al., 2006; Shah et al., 2011): fast Na^+^ (NaF), delayed rectifier K^+^ (KDR), *A*-type K^+^ (KA), *L*-type Ca^2+^ (CaL), calcium gated K^+^ (SK), hyperpolarization activated cyclic nucleotide gated (HCN), *M*-type K^+^ (KM) and *T*-type Ca^2+^ (CaT) channels. The model received place field inputs through 100 different conductance-based synapses (Narayanan and Johnston, 2010), whose independent afferent inputs during place-field traversals were stochastic and governed by a Gaussian-modulated cosinusoidal (8 Hz) distribution (Fig. 1*C*). The resulting intracellular voltage response of a model endowed with such synaptic afferents was reflective of theta-modulated firing (Fig. 1*D*) observed during place-field traversals (Harvey et al., 2009; Basak and Narayanan, 2018).

To obtain the phase code of space corresponding to a model with specific ion channel conductances that define its intrinsic properties, we computed the firing profiles of the model neuron for 50 consecutive one-dimensional place-field traversals (Figs. 1–2). These consecutive inputs were fed to the model neuron through their synapses, with their place field centers separated by *T* ms, the time taken for the virtual animal to traverse between these centers (Fig. 1*E*). Additionally, consistent with the TSC framework, the theta phase associated with every place-field input was shifted by *τ* ms (Fig. 1*E*) with reference to its immediately preceding place-field input (Skaggs et al., 1996; Geisler et al., 2010). The firing profiles corresponding to these 50 consecutive place-field traversals were combined to construct a population firing rate (*f*_POP_) profile (Fig. 1*F*), which was considered as the extracellular reference theta (Geisler et al., 2010). As expected from the TSC framework, the frequency of this reference theta was lesser (Fig. 1*F*, inset; 7.57 Hz) than the intracellular theta (8 Hz) conforming to the relationship between intracellular and reference theta waveforms in the hippocampus (Harvey et al., 2009; Geisler et al., 2010). Spike times corresponding to each of the 50 traversals were then temporally aligned with this reference theta to compute the associated spike phases (Geisler et al., 2010), by defining individual oscillatory cycles to cover consecutive troughs in the reference theta (Fig. 1*F*). This phase computation demonstrated that our model exhibited precession of phase within each traversal (Fig. 1*G*), thereby providing a conductance-based model for evaluating phase code of space within the TSC framework.

### Neuronal intrinsic properties play a critical role in defining the phase code and its efficiency in encoding the space within a place field

Phase precession constitutes a phase code of space in the external world. How efficient is this phase code in transferring information about space (spanning a single place field) encoded by a place cell? Do neuronal intrinsic properties play a role in regulating efficiency of information transfer through such phase codes? Motivated by the efficient coding literature that recruits maximization of information transfer as a quantitative metric (Barlow, 1961; Bell and Sejnowski, 1995, 1997; Simoncelli and Olshausen, 2001; Lewicki, 2002; Simoncelli, 2003), we defined the efficiency of this phase code based on maximal mutual information between the stimulus and response, considering space to be the stimulus and spike phase to be the response. This representation, which did not place any parametric constraints on the nature of phase precession, allowed for a generalized definition of coding efficiency.

To assess the implications of neuronal intrinsic properties on the phase code and its efficiency, we computed phase coding efficiency in models that received *identical* inputs during place field traversals, and were distinct in terms of the channel densities that defined them. Methodologically, as a virtual animal traversed across 50 successive placed fields, the temporal structure of the afferent drive onto individual model neurons, each with disparate intrinsic properties, was set to be *identical*. Specifically, models were distinct from each other only in terms of their intrinsic properties (parameters in Table 1), with no differences in the temporal structure of afferent inputs. Constructing and comparing the spike-phase profiles of such models provided us the first clue on the strong dependence of spike phase precession on neuronal intrinsic properties. Two extreme examples elucidating this point are shown in Fig. 3*A–B*, with the model in Fig. 3*A* eliciting bursts of spikes throughout the entire place field, resulting in a code where the phases assigned to individual spatial bins within the place field (Fig. 3*A*, left) were not delineated (Fig. 3*A*, right). These overlaps in phase responses to spatial stimuli implied that this phase profile carried little information about the spatial location of the animal (MI=0.04 bits).

Although both models received identical afferent inputs and had their phase codes computed with precisely the same procedure under the TSC framework, only the model in Fig. 3*B* manifested a phase code that exhibited phase precession, with a clear monotonic relationship to spatial location. This translated to broadly well-delineated range of phases being assigned to specific spatial bins (Fig. 3*B*, right), implying that the spatial information contained in this phase code was higher (Fig. 3*B*; MI=2.1 bits) than the model in Fig. 3*A*. The differences between these two models were limited to intrinsic properties, and not in the temporal structure of afferent network inputs or associated *T*/***τ*** interactions (Fig. 1*E*) that define phase precession within the TSC framework. Therefore, these observations pointed to a pivotal role for neuronal intrinsic properties in the emergence of phase precession and in regulating the efficiency of the associated phase code.

### Degeneracy in efficient phase coding

Is such dependence of the phase code on neuronal intrinsic properties extremely constrained, whereby phase precession emerges only for a small cluster within the model parametric space? We reasoned that hand-tuning of channel properties to obtain one specific model and exploring parametric dependencies in that single model entails biases that would make our conclusions to be mere reflections of those specific parametric choices. Therefore, we implemented an unbiased stochastic search approach (Foster et al., 1993; Prinz et al., 2004; Marder and Goaillard, 2006; Marder, 2011; Marder and Taylor, 2011; Rathour and Narayanan, 2012, 2014; Anirudhan and Narayanan, 2015; Mukunda and Narayanan, 2017; Basak and Narayanan, 2018; Mittal and Narayanan, 2018; Mishra and Narayanan, 2019), where we built several models with distinct parametric combinations to assess the phase-space dependence with a heterogeneous population of neuronal models. We designed this parametric search space to be wide (Table 1) to avoid biases in parametric choices and to incorporate heterogeneities spanning neuronal biophysical properties (Foster et al., 1993; Prinz et al., 2004; Marder and Goaillard, 2006; Marder, 2011; Marder and Taylor, 2011; Rathour and Narayanan, 2012, 2014; Anirudhan and Narayanan, 2015; Mukunda and Narayanan, 2017; Basak and Narayanan, 2018; Mittal and Narayanan, 2018; Mishra and Narayanan, 2019; Rathour and Narayanan, 2019).

In implementing the stochastic search, we picked each of the 11 different parameters from their respective uniform distributions (Table 1) and constructed 11,000 unique model neurons. We computed the responses of each of these models to all 50 place-field traversals and computed *f*_POP_ as the reference theta oscillation for each model. We calculated spike phases with reference to *f*_POP_, and constructed the multi-traversal phase-space profile for each model (Fig. 2). We computed mutual information on this phase-space profile to assess the dependence of phase code efficiency on neuronal intrinsic properties of these models. Although the temporal structures of their afferent inputs were identical, these models showed a wide range of mutual information values, confirming that the efficiency of the phase code was critically dependent on neuronal intrinsic properties.

What were the constraints on the underlying parameters in models that achieved high phase code efficiency? Do such efficient models manifest clustering of underlying parameters, suggesting the requirement of a unique parametric regime for efficient phase coding? To address these questions, we first picked five (of the 11,000) models that were endowed with very similar phase precession profiles with high efficiency in spatial information transfer (Fig. 4*A*). When we plotted the parameters associated with these five models (Fig. 4*B*), we found a lack of any clustering in the parametric combinations that resulted in these highly efficient models with very similar phase precession. This suggested the expression of degeneracy (Edelman and Gally, 2001) in the manifestation of efficient phase codes, where disparate parametric combinations (representing distinct ion channels) yielded similar function (phase-coding efficiency).

To further evaluate degeneracy in efficient model populations, we picked 284 high-efficiency models among the 11,000 models by setting a cut-off of 1.5 bits (Fig. 3*C*) on the space-to-phase mutual information (Fig. 4*D*). We explored potential clustering in parameters by plotting the histogram for each of the 11 parameters associated with these 284 efficient models with MI ≥ 1.5 bits (Fig. 4*C*, bottom row). The broad distribution of parametric values that yielded these efficient models provided clear evidence for the expression of degeneracy in the emergence of efficient phase coding. Could the emergence of these efficient models be dependent on correlated expression of ion channel conductances? To assess this, we plotted pair-wise scatters of the 11 parameters from all the 284 models (Fig. 4*C*), and computed the associated correlation coefficients (Fig. 4*E*). We found the pairwise correlations among model parameters to be weak, with the Pearson’s correlation coefficient spanning the range of –0.5 ≤ *R* ≤ 0.5 (Fig. 4*E–F*). In summary, although neuronal intrinsic properties played a critical role in the emergence of phase precession and in regulating the associated efficiency, there were several nonunique parametric combinations with weak pair-wise correlations that yielded similar high-efficiency models.

### Degeneracy in concomitant emergence of efficient phase coding and robust intrinsic excitability

The analyses thus far did not account for the characteristic intrinsic excitability of CA1 pyramidal neurons. Do models that are efficient in terms of encoding space through phase also exhibit signature excitability characteristics of the specific neuronal subtype? To address this, we picked five measurements (resting membrane potential, RMP; standard deviation of RMP; input resistance; firing rates for pulse current injections of 50 pA and 250 pA) that govern CA1 pyramidal neuron excitability, and asked if the 284 efficient models also satisfied electrophysiological bounds (Narayanan and Johnston, 2007; Narayanan et al., 2010; Malik et al., 2016; Rathour et al., 2016; Das and Narayanan, 2017) on these measurements (Table 2). This additional validation process resulted in 132 models that were efficient, and *concomitantly* satisfied multiple constraints on signature intrinsic excitability characteristics.

Did the imposition of an additional layer of excitability constraints weaken the degeneracy that was expressed when efficient phase coding was the only validation criterion (Fig. 4)? To address this question, we first picked five example models that were concomitantly efficient and endowed with signature excitability. These models had similar or identical values not only for the mutual information measure (Fig. 5*A*) but also for each of the 5 intrinsic measurements (Fig. 5*B–D*). Despite similarities in phase-coding efficiency and in the five intrinsic measurements, the underlying parametric values that defined these five models exhibited a broad distribution (Fig. 5*E*). We analyzed the distributions of phase code efficiency (Fig. 6*A*) and of the different measurements of intrinsic excitability (Fig. 6*B–D*), encompassing all the 132 valid models that were concomitantly efficient and had signature excitability characteristics. The distributions of the electrophysiological measurements demonstrated the heterogeneities inherent to these models, where a tight clustering in these measurements was absent. Consistent with our observations with the five example models (Fig. 5), we found that the parametric distributions of all 11 parameters spanned a broad range rather than showing specific clusters (Fig. 6*E*; last row histograms). We also noted that there were no strong pairwise correlations across all parametric combinations (Fig. 6*E–G*).

Together these results showed that neither the requirement on high efficiency of the phase code nor the multiple additional validation criteria on signature excitability properties were sufficient to enforce strong constraints on the distributions of parameters. These observations point to the existence of disparate routes to achieve similar phase coding efficiency and neuronal excitability. The existence of such disparate routes unveils degeneracy in the concomitant emergence of information-rich encoding and homeostasis in neuronal excitability.

### Synergistic functional interactions between synaptic strength and intrinsic excitability governed the emergence of efficient phase codes and associated parametric degeneracy

Although the model parameters did not exhibit strong correlations in efficient models, was there any correlation at the functional level between the efficiency of the phase code and the characteristics of the rate code? To address this, we plotted phase-coding mutual information against the maximal firing frequency *F*_max_ (Fig. 7*A*) or full-width at half maximum (FWHM) of the firing rate profile (Fig. 7*B*) of the model cell within the place field. We found that neurons with high phase coding MI were typically obtained when the peak firing rate was around 5 Hz (Fig. 7*A*) and when the FWHM was around 1 s (Fig. 7*B*). As FWHM is a function of the dorso-ventral location of the pyramidal neuron (Kjelstrup et al., 2008; Strange et al., 2014) and is a relative (to the spread of the input distribution) measure within our modeling framework, it would be infeasible to directly extrapolate the observations here to electrophysiological. However, we found that firing rates where high phase-coding MI was achieved (Fig. 7*A*) were comparable to *in vivo* recordings from CA1 place cells (Harvey et al., 2009; Cohen et al., 2017).

Was there any correlation at the functional level between the efficiency of the phase code and synaptic/intrinsic properties? Was the emergence of efficient phase codes dependent on correlations between synaptic and intrinsic measurements? First, we computed correlations between the phase-coding efficiency of the models (*i.e.*, MI values) and each of the three excitability measurements, separately for the 284 efficient models and the 132 efficient and excitability-validated models (Fig. 7*C–E*). We found these correlations to be extremely weak (Fig. 7*C–E*) thereby ruling out a direct well-defined relationship between intrinsic excitability and the model’s ability to efficiently encode space through phase.

Second, as the afferent drive was identical across all models, we employed synaptic permeability (*P*_max_ in Eq. 5, a measure of receptor density) in individual models as the functional equivalent for synaptic strength. With this equivalence, we found the correlation between model efficiency and synaptic strength for the two groups of valid models to be weak (Fig. 7*F*). Although there was no strong correlation at the functional level between the efficiency of the model and its intrinsic/synaptic properties, we found strong negative correlations between neuronal intrinsic properties and synaptic permeability for both valid model populations (Fig. 7*G–I*). This negative correlation was particularly strong between neuronal firing rate and the synaptic permeability (Fig. 7*I*), suggesting that phase-coding efficiency and associated parametric degeneracy were mediated by the synergy between intrinsic and synaptic properties. Specifically, the ability of intrinsic excitability and synaptic drive to *counterbalance* each other played a critical role in defining models with high-efficiency phase codes and in driving associated parametric degeneracy.

Thus far in our analyses, we first sorted models on the basis of their efficiency (Fig. 3–4) and among highly efficient models found a subset that was also endowed with signature excitability characteristics (Fig. 5–7). Instead, if models were initially sorted by whether they were endowed with signature excitability characteristics irrespective of what their MI values were, would these intrinsically valid models also be efficient phase coders? Would there be significant correlations between intrinsic properties and MI in this intrinsically valid model population? To address these, we identified models (Fig. 8*A*; 1,754 out of 11,000) that were intrinsically valid (Table 2) irrespective of what their MI values were (Fig. 8). We found the MI values of these intrinsically valid models to span the 0–2 bits range (Fig. 8*B*), implying that they were not necessarily efficient and providing further evidence that efficiency in models was not a simple reflection of signature neuronal excitability properties. Additionally, we confirmed that our earlier (Fig. 7) conclusions on weak correlations between MI and intrinsic/synaptic properties extended to these intrinsically valid models as well (8*B–E*). Importantly, we found that the synergy between synaptic and intrinsic properties, manifesting as high correlations between intrinsic measurements and synaptic permeability, was observed only in model populations that were endowed with high-efficiency phase codes (Fig. 7*G–I*), but was notably absent in these models that were just intrinsically valid (Fig. 8*F–H*). These observations demonstrated that the synergy between intrinsic and synaptic properties was not a reflection of the stochastic search process, but manifested as an essential cog in the emergence of efficient models and associated degeneracy.

### The impact of virtually knocking out individual channels on efficient phase coding was differential and variable

Although we had established the expression of degeneracy in the emergence of efficient phase codes, assessment of the impact of individual channels provides insights and predictions about their dominance hierarchy in establishing such emergence. Specifically, although several channels could contribute and regulate a specific physiological property, there are specific channels whose contribution to the property is dominant (*e.g.*, the role of sodium channels in action potential generation). Within our conductance-based framework for phase precession, are there specific sets of ion channels that support efficient transfer of spatial information through spike phase?

An ideal way to assess the relative roles of individual channels within the degeneracy framework is the virtual knockout strategy (Rathour and Narayanan, 2014; Anirudhan and Narayanan, 2015; Mukunda and Narayanan, 2017; Basak and Narayanan, 2018; Mittal and Narayanan, 2018). In adapting this strategy for assessing the impact of individual channels on efficient phase coding, we virtually knocked out individual channels in our models by setting the corresponding conductance to zero, with no changes to any other parameter or inputs to the selected model. After such knockout of individual channels, we computed the multi-trial phase-space plot of each virtual knockout model (VKM). We compared the mutual information computed from this phase-space plot with that from the model when the channel conductance was intact, and calculated the percentage change in mutual information after virtual knockout of the specific channel (Fig. 9*A*). We repeated this procedure for each of the 132 valid models, and individually for the 6 active channel subtypes (Fig. 9; 132 × 6 = 792 VKMs, each subjected to 50 traversals). We did not perform virtual knockout analyses on the NaF and KDR channels, because these VKMs ceased spiking upon knocking out either of these channels, implying that spike phases could not be computed.

The impact of knocking out each of the 6 channels on the phase-space plot is illustrated for an example model (Fig. 9*A*). The distribution of percentage changes in the mutual information in the population of models after virtual knockout of each of the 6 individual channels (Fig. 9*B–C*) unveiled the heterogeneity of the impact of these channels on efficient phase coding. Specifically, the impact of knocking out each of these channels was variable, whereby there was a strong effect on MI in some models with others showing no significant effect upon knockout of the same channel subtype. From the cellular perspective, the impact of knocking out different channels had differential effects on MI, with considerable cross-cellular heterogeneity in the relative contributions of individual channels to MI (Fig. 9*B–C*).

We asked if the impact of one channel subtype on efficient phase coding could predict the impact of another channel, by comparing percentage changes in MI after individual channel knockouts in a pairwise manner (Fig. 9*D*). Consistent with the correlation analyses on channel conductances (Fig. 6*E–G*), we found that the pair-wise correlations between percentage changes in MI after channel knockouts were very weak (Fig. 9*D–F*). The many-to-one relationship between different channels and coding efficiency, derived from variability in and weak correlations among the effects of individual channels, forms the substrate for ion channel degeneracy in the emergence of efficient phase codes (Fig. 4–6). From the perspective of dominance of individual channels, the statistics of our VKM analyses present a testable prediction on a critical role of SK and *T*-type calcium channels in determining the efficiency of the phase code (Fig. 9*B–C*).

### Asymmetry in place-field afferent inputs introduces predictive temporal shifts to the rate and phase codes with the adaptive temporal shift preserving phase-code efficiency

There are theoretical and experimental lines of evidence for an experience-dependent asymmetric expansion of hippocampal place fields in the direction opposite to the movement of the animal (Mehta et al., 1997; Mehta et al., 2000; Mehta et al., 2002; Harvey et al., 2009). However, the place-field model employed here involves a Gaussian that is symmetric with reference to the place cell center (Fig. 1*C*). Phase precession within the TSC framework is attributed to the specific temporal relationships at the behavioral scale (*T*) and the theta-time scale (***τ***) (Harvey et al., 2009; Geisler et al., 2010). Therefore, the symmetric nature of the afferent input activation was irrelevant in the emergence of phase precession (Figs. 1–2) within this framework (Geisler et al., 2010). What is the impact of asymmetry in the profile of afferent input activation on the place-cell rate and phase codes within the TSC framework? Does such a change in the statistics of place field-driven afferent inputs onto a single neuron alter the efficiency of the phase code in the model? Does the phase code adapt to such changes in afferent statistics to preserve efficiency?

We replaced the symmetric Gaussian profile of afferent synaptic activation (Fig. 1*C*) by a horizontally reflected Erlang distribution while matching the area under the curve of these profiles (Fig. 10*A*). Similar to the theta modulation that we had introduced for the Gaussian, we incorporated theta modulation to the Erlang distribution by multiplying this function with an 8-Hz sinusoid. We maintained the same *T*-***τ*** relationship as that of Gaussian profile of synaptic activation (Fig. 1*E*; Fig. 2) for the 50 consecutive place-field traversals, which now involved the Erlang distribution instead. We activated all our 132 models that were efficient and concomitantly matched CA1 excitability properties (Fig. 6) to 50 place field traversals and computed the firing rates of these model cells within the place fields (Fig. 10*B*). We repeated this procedure for three different values (40 Hz, 80 Hz and 160 Hz) of the maximal afferent activation rate (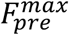; Eq. 7, Eq. 19; see Fig. 10*A*). Predictably, the firing rate of the model cells increased with increase in 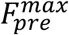(Fig. 10*B–C*), irrespective of whether the activation profile was driven by a symmetric or an asymmetric distribution. As would be expected from the asymmetry in the afferent activation profile, we also found that the rate code displayed an asymmetry when the models were activated with the asymmetric distribution, with asymmetry progressively increasing with increase in 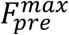(Fig. 10*B*, Fig. 10*D*).

How did the phase code change as functions of the asymmetry and 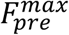? To address this, we computed the phase-space plots for each of these models with each input configurations (symmetric *vs*. asymmetric and three different values of 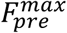) by computing the spike phases with reference to the respective *f*_POP_. We found phase precession to be overall similar for symmetric *vs*. asymmetric synaptic activation profiles (Fig. 10*E–F*), which was confirmed by the similar efficiency in the phase code (Fig. 10*G*). An important effect of asymmetry in the afferent activation profile was a leftward “predictive” shift in the phase precession profile, which was consequent to the early intra-place-field firing when neurons were activated with an asymmetric input profile and was observed for all values of 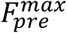 (Fig. 10*B*, Fig. 10*D*).

These analyses revealed important differences in the phase code that emerged when the synaptic activation rate 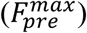 was changed (Fig. 10*E–G*). First, consistent with the rate code (Fig. 10*B*), the temporal spread of the extent of place-field firing increased with 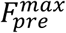, implying that the phase code now spread over a larger number of theta cycles (Fig. 10*E*). Second, the increase in firing rate with increase in 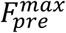 implied that the cell spiked more than once during a single theta cycle (Fig. 10*E–F*), resulting in a reduction in the efficiency of the phase code with increase in 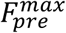, when all spikes were considered for the computation of mutual information (Fig. 10*G*). Third, changes introduced to the phase code by increasing 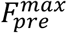 were broadly invariant to whether the input activation profile was symmetric or not. Specifically, the increase in temporal extent of firing, the presence of multiple spikes within a single theta cycle and the reduction in the efficiency of the phase code were all observed with increase in 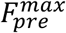, irrespective of whether the synaptic activation profile was symmetric or asymmetric (Fig. 10*E–G*). Together, in our model, asymmetry in place-field afferent inputs introduced predictive temporal shifts to the rate and phase codes, with the shift in the phase code constituting a stimulus-dependent adaptation of the code to the altered input statistics in order to achieve similar efficiency. These results imply that the phase code follows the stimulus statistics, which in this case is driven by the distribution of the afferent inputs as a function of space (as the traversal itself was considered uniform), thereby preserving the efficiency of information transfer.

### Theta modulation and symmetry profiles of sub-threshold ramps in place-cell voltage responses

Thus far, we have shown that our model neurons exhibit signature electrophysiological and encoding characteristics of CA1 place fields. Specifically, we showed that our place cell models exhibit electrophysiologically-matched intrinsic properties (Figs. 5–7), also manifesting biological heterogeneities in these intrinsic properties (Rathour and Narayanan, 2012, 2014; Cembrowski et al., 2016; Malik et al., 2016) through variable expression of ion channels expressed in these neurons. Importantly, the kinetics and gating properties of these ion channels and the range of intrinsic heterogeneities were derived from CA1 pyramidal neuron electrophysiology. The incorporation of these intrinsic heterogeneities provided the substrate for us to demonstrate the need for strong synergy between intrinsic and synaptic properties to achieve efficient phase-coding models (Fig. 7*I*). We showed that our models matched place-field firing rates (Fig. 1*D*; Fig. 7A–B; Fig. 10) and experience-dependent asymmetry (Mehta et al., 1997; Mehta et al., 2000; Mehta et al., 2002; Harvey et al., 2009) in their firing rate profiles (Fig. 10). These place-cell electrophysiological characteristics that our models matched were in addition to the core focus of our study on achieving phase precession within the TSC framework (Skaggs et al., 1996; Harvey et al., 2009; Geisler et al., 2010), where we had achieved phase precession by emergence of the extracellular theta with a frequency that was *lesser* than the intracellular theta frequency (Fig. 1F–G). In addition to these characteristics, place cells exhibit strong theta-power and manifest a sub-threshold ramp in their voltage response during place-field traversal (Harvey et al., 2009). To further validate our models against these characteristics, we assessed theta power and sub-threshold ramps in our model voltage responses and asked if they matched with their electrophysiological counterparts. We performed these for each of the 132 models, with 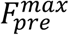 set to three different values and with both symmetric as well as asymmetric synaptic activation profiles (Fig. 11).

We analyzed theta modulation and theta power by band-pass filtering the voltage response between 4–10 Hz. The sub-threshold voltage response profile, to verify the presence of and quantify sub-threshold ramps, was computed by median filtering the voltage response trace. The raw voltage responses, the theta-filtered traces and the sub-threshold response profiles for an example model neuron, receiving symmetric (Fig. 11*A*) and asymmetric (Fig. 11*B*) synaptic activation profiles for the three different values of 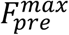 (40, 80 and 160 Hz) confirmed that our models manifested signature electrophysiological characteristics. Specifically, we observed that the theta-filtered traces showed clear increase in theta power in the voltage response during place-field traversals, with the spectra consistently showing peak power at 8 Hz. We also noted a consistent sub-threshold response profile of 5–8 mV amplitude that was either symmetric (Fig. 11*A*) or exhibited an asymmetric ramp (Fig. 11*B*) depending on the nature of synaptic activation. We noted that both the theta modulation (amplitude of the band-pass filtered signal) and the membrane potential ramp amplitude are comparable to those observed in place cells *in vivo* (Harvey et al., 2009).

We quantified these measurements for all the 132 models that are both information-efficient and intrinsically valid, for all the six cases (three different values of 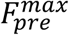 for symmetric and asymmetric activation profiles) of synaptic activation profiles. We computed the area under the curve (AUC) of the Fourier spectrum of the theta-filtered voltage response (Fig. 11*C*), the ramp amplitude (Fig. 11*D*) and asymmetry index (Eq. 21; Fig. 11*E*) from the correspondingly filtered versions of these voltage responses. Our results confirmed the expression of theta modulation and high theta power during place-field traversals across all models (Fig. 11*C*). We also confirmed that there was always a depolarizing sub-threshold voltage profile during place field traversals that formed the substrate for action potential firing (Fig. 11*D*). This sub-threshold voltage profile was an asymmetric ramp (Fig. 11*B*) showing a strong asymmetry index (Eq. 21) when the model was presented with asymmetric synaptic activation profiles (Fig. 11E). Together, our model measurements matched their electrophysiological counterparts with reference to the expression of strong theta modulation and the manifestation of a depolarizing sub-threshold ramp during place-field traversals.

## DISCUSSION

We showed that phase coding in place cells is critically reliant on a synergistic balance between intrinsic and synaptic properties, and that efficient information transfer through such a phase code could be achieved through multiple disparate routes while concomitantly maintaining signature excitability properties. We demonstrated these by developing a conductance-based model for phase precession within the TSC framework, and by defining efficiency of the phase code by recruiting the information maximization formulation. In adapting the TSC framework to a conductance-based setting, we performed an unbiased stochastic search across parameters involving thousands of models to ensure that we capture biological heterogeneities. This stochastic search process yielded models that matched several signature sub- and supra-threshold electrophysiological characteristics of CA1 place cells, and unveiled degeneracy in the concomitant emergence of efficient phase coding and robust excitability characteristics. We showed that the impact of individual ion channels on phase-code efficiency was differential and variable, with an experimentally testable prediction on the critical role of SK channels in governing phase precession and associated efficiency. Finally, by modifying the symmetry properties of the afferent inputs within the TSC framework, we demonstrated that asymmetry in place-field afferent inputs introduces predictive temporal shifts to the rate and phase codes. We noted that the shift in the phase code constitutes an adaptive shift to preserve phase-code efficiency in a manner that was driven by afferent stimulus statistics.

### Synergistic interactions between intrinsic and synaptic properties drive phase coding efficiency

Our results make a clear case for phase precession and the efficiency of the associated phase code to be regulated by neuronal intrinsic properties, rather than being solely reliant on the temporal structure of the afferent network inputs. Employing models that *received afferent inputs with identical temporal structure*, we showed that neuronal intrinsic properties are critical in achieving efficient phase coding. Importantly, this dependence was not driven by a simple correlation between efficient phase coding and neuronal excitability (Fig. 7–8). Instead, phase-coding efficiency and associated parametric degeneracy were mediated by synergistic interactions between intrinsic and synaptic properties, specifically pointing towards the ability of intrinsic excitability and synaptic drive to *counterbalance* each other in achieving this (Fig. 7*I*). Based on these observations, we postulate that the emergence of stable, efficient and robust encoding in neuronal systems relies on synergistic interactions between disparate forms of plasticity. Under such a postulate, the specific forms of plasticity that define such emergence would be variable in a neuron- and context-dependent manner, depending on the internal state of the network (given parametric degeneracy) and on the afferent modulation imposed by behavior. Future studies could explore the manifestation of such counterbalances in intrinsic *vs*. synaptic characteristics, and their roles in regulating encoding *and* homeostasis under physiological or pathophysiological conditions where these characteristics are known to undergo changes.

Our model presents a quantitative testable prediction on the dominant impact of SK channels on phase coding. Future computational studies could focus on whether this dominant role is a reflection of the slow kinetics of the SK current, or if this were just a reflection of the SK current altering neuronal excitability. Experimentally, the role of SK currents on phase coding in place cells could be tested with pharmacological agents or transgenic mice that have altered SK channel conductance or properties.

### Degeneracy in efficient coding and excitability robustness

We show that the emergence of efficient phase coding (Figs. 4) and *concomitant* excitability robustness (Fig. 5–6) is independent of the ability of a neuron to maintain its ion channel densities at *specific* values, but was driven by synergistic interactions between synaptic and intrinsic properties (Fig. 7*I*). Such degeneracy implies that there are several degrees of freedom available to a neuron in concomitantly maintaining efficiency of the phase code *and* homeostasis of intrinsic excitability, without cross interferences between the encoding and the homeostasis processes. This constitutes an important departure from conventional analyses of the encoding-homeostasis balance, where encoding is hypothesized to be achieved by *specific* processes and *other* concurrent (or slower) processes achieve homeostasis. Within our framework, encoding and homeostasis is postulated to emerge concomitantly, with significant *degeneracy* in the specific components that contribute to such emergence.

Our analyses also constitute a scenario where redundancy reduction with reference to a code is brought about by degeneracy in the structural components that contribute to the emergence of the code. It is important to note that our analysis does not constitute coding degeneracy, where disparate codes (potentially mediated by different structural components) encode the same stimulus or structural redundancy, where a dysfunctional component is replaced by an identical component that restores function. Ours is an example of a scenario where an efficient code, that reduces redundant representations, is achieved by disparate combinations of underlying structural components (channels and receptors). Similar to other examples of degeneracy across the literature (Foster et al., 1993; Edelman and Gally, 2001; Prinz et al., 2004; Marder, 2011; Basak and Narayanan, 2018; Rathour and Narayanan, 2019), while the contribution of different structural components to individual models is variable, the specific function that emerges as a consequence of interactions between these distinct structural components remains *precise and well defined*.

### Limitations of our model and future directions

While emphasizing on the phase code from a single neuron perspective, our approach did not account for the rate code or for phase and rate codes from a network perspective where different neurons together encode space by representing different place fields in an arena (Mehta et al., 2002; Huxter et al., 2003; O’Keefe and Burgess, 2005). The analyses of encoding efficiency that accounts for multi-neuronal rate and phase coding is an important step forward. Such analyses should assess the specific roles of neuronal intrinsic properties in the concomitant emergence of efficient rate and phase codes across neurons, along with efficacious maintenance of intrinsic neuronal excitability across the network. These analyses would also provide avenues for assessment of degeneracy in a network-coding framework, for the analysis of the role heterogeneities in place field properties (*e.g.*, in peak firing rates of individual neurons, extent of individual place fields, and the slope of phase precession), and for the study of potential relationships between phase-coding efficiency at a single-neuron scale and network-scale, spanning a large spatial arena.

In the context of experience-dependent asymmetry, our model predicts a temporal adaptation that preserves the efficiency of the phase code when the symmetry of afferent synaptic drive is altered (Fig. 10). Our results show an expected predictive temporal shift in the rate and phase codes, with a stimulus statistics-dependent adaptation that preserved phase-coding efficiency (Fig. 10). However, it is important to note that the asymmetric afferent drive is just one of the physiological attributes that change with experience, with other attributes such as the somatodendritic inhibitory tone (Sheffield et al., 2017), the overall afferent drive and dendritically initiated spiking (Cohen et al., 2017) also exhibiting changes with experience. Although we report stimulus-dependent adaptation in the phase code that preserved its efficiency with the asymmetry, experience-dependence might alter or preserve the efficiency of the phase code through any of these other experience-dependent changes, which have not been incorporated into our model. Therefore, future models could assess experience-dependence of phase coding efficiency (and potential mechanisms that preserve efficiency) by accounting for all aspects of experience dependence, rather than assessing only asymmetric afferent drives (Mehta et al., 1997; Mehta et al., 2000; Mehta et al., 2002; Cohen et al., 2017; Sheffield et al., 2017).

Accounting for these aspects of experience dependence also requires that the place-cell model accounts for morphological properties of CA1 pyramidal neurons (Basak and Narayanan, 2018), including localization and activation profiles of intrinsic (Rathour and Narayanan, 2014; Basak and Narayanan, 2018; Hsu et al., 2018) and synaptic (Klausberger and Somogyi, 2008; Sheffield and Dombeck, 2015; Sinha and Narayanan, 2015; Grienberger et al., 2017; Basak and Narayanan, 2018; Boivin and Nedivi, 2018) properties. Incorporation of these components into morphologically realistic neurons will enable the computation of extracellular theta waveforms using forward modeling approaches for local field potentials (Einevoll et al., 2013; Sinha and Narayanan, 2015), rather than using population firing as a proxy for the extracellular theta. We postulate that the degeneracy in efficient phase coding and concomitant robustness in intrinsic excitability, emergent due to synergistic interactions between intrinsic and synaptic properties, would span more structural components and express more effectively if these additional components and associated interactions were introduced into our model.

Finally, our model was limited to phase precession in CA1 pyramidal neurons, focusing on heterogeneities in intrinsic properties owing to baseline variability, activity-dependent plasticity or neuromodulation. We postulate that our fundamental conclusions would extend to other neuronal subtypes that exhibit phase precession. Specifically, we hypothesize that synergistic interactions between intrinsic and synaptic properties would mediate the expression of degeneracy in the concomitant emergence of efficient phase coding and homeostasis in other neuronal subtypes as well. Such degeneracy would be achieved through disparate combinations of *specific* structural components expressed in those neuronal subtypes, and should be explored through careful assessment of intrinsic–synaptic counterbalances mediated by ion channels and receptors expressed in those neurons.

## Acknowledgments

This work was supported by the Wellcome Trust-DBT India Alliance (Senior fellowship to RN; IA/S/16/2/502727), Human Frontier Science Program (HFSP) Organization (RN), the Department of Biotechnology (RN), the Department of Science and Technology (RN), and the Ministry of Human Resource Development (RN & PS). The authors thank the members of the cellular neurophysiology laboratory for helpful discussions and for comments on a draft of this manuscript.

